# Allosteric modulators enhance agonist efficacy by increasing the residence time of a GPCR in the active state

**DOI:** 10.1101/2021.01.05.424557

**Authors:** Anne-Marinette Cao, Robert B. Quast, Fataneh Fatemi, Philippe Rondard, Jean-Philippe Pin, Emmanuel Margeat

## Abstract

Much hope in drug development comes from the discovery of positive allosteric modulators (PAM) that display target subtype selectivity, and act by increasing agonist potency and efficacy. How such compounds can allosterically influence agonist action remains unclear. Metabotropic glutamate receptors (mGlu) are G protein-coupled receptors that represent promising targets for brain diseases, and for which PAMs acting in the transmembrane domain have been developed. Here, we explore the effect of a PAM on the structural dynamics of mGlu2 in optimized detergent micelles using single molecule FRET at submillisecond timescales. We show that glutamate only partially stabilizes the extracellular domains in the active state. Full activation is only observed in the presence of a PAM or the G_i_ protein. Our results provide important insights on the role of allosteric modulators in mGlu activation, by stabilizing the active state of a receptor that is otherwise rapidly oscillating between active and inactive states.

## Introduction

G protein-coupled receptors (GPCR) constitute the largest family of integral membrane receptors encoded in the human genome and are involved in various physiological processes (Lagerström and Schiöth 2008). They constitute the main targets in drug development programs for many therapeutic applications (Alexander et al. 2019). Within recent years, much hope came from the discovery of allosteric modulators targeting GPCRs with a few already on the market (May et al. 2007; Conn et al. 2009; Thal et al. 2018; Changeux and Christopoulos 2016). Their pharmacological interest comes from their ability to target allosteric sites different from the evolutionary conserved orthosteric site, conferring higher subtypes selectivity. Most importantly, positive allosteric modulators (PAM) enhance agonists effects on GPCR, then preserving their rhythm of biological activity where and when needed physiologically (Foster and Conn 2017). PAMs can display various effects (Christopoulos 2014), including increasing agonist potency (from 2 to 100 fold), increasing agonist efficacy, partially activating receptors (ago-PAM effect) or even orienting the receptor towards one of its signaling pathway (Rook et al. 2013; Makita et al. 2007). It is commonly considered that PAMs act by stabilizing a specific conformation of the receptor (Shaye et al. 2020; Bueno et al. 2020; Kruse et al. 2013; Liu et al. 2019; Srivastava et al. 2014). However, PAMs may likely act by influencing the equilibrium between preexisting GPCR conformational states.

Class C GPCRs are especially amenable for allosteric modulation, notably given their highly modular architecture, being more complex than the simple rhodopsin-like structure (Pin and Bettler 2016). These receptors include the metabotropic glutamate (mGlu), the GABA (GABABR), the calcium-sensing (CaSR), and the umami and sweet taste receptors (T1R) (Møller et al. 2017). The mGlu receptors are responsible for the modulatory activity to L-glutamate (Glu), the major excitatory neurotransmitter in the central nervous system, and are therefore essential in the fine-tuning of synapses (Gregory and Goudet 2021). Class C GPCRs are composed of two subunits, each comprising several functional domains **(Fig. 1)**. The large extracellular domain (ECD) consists of a Venus flytrap domain (VFT), harboring the orthosteric site, and a rigid linker connected to the 7 transmembrane domain (7TM) (Koehl et al. 2019; Shaye et al. 2020; Zhang et al. 2020). Most identified class C GPCR allosteric modulators act in the 7TM at a site corresponding to the orthosteric site of the rhodopsin-like GPCRs (**Fig. 1**) (Doré et al. 2014; Wu et al. 2014; Christopher et al. 2018; Rovira et al. 2015). Other sites have also been identified close to the orthosteric site (Servant et al. 2010; Zhang et al. 2008), at the active interface of the VFT dimer (Scholler et al. 2017), or at the active interface of the 7TM dimer (Shaye et al. 2020). Despite our knowledge of their binding mode, how such molecules allosterically control agonist affinity or efficacy, exert partial agonist activity or biased effect remains largely unknown.

**Figure 1:**
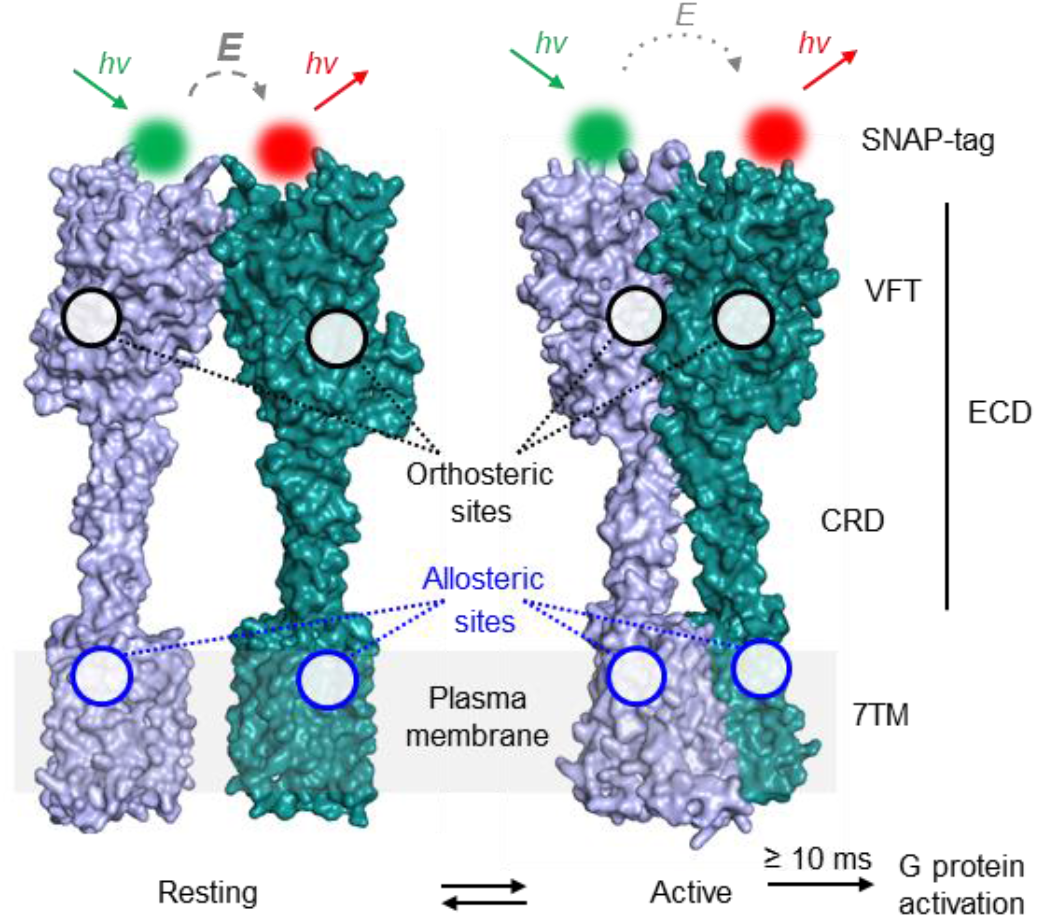
Structure and conformational rearrangements of mGlu receptor. Structural model of dimeric mGlu2 in resting and active conformations. The major structural elements of each subunit include the extracellular domain (ECD)(comprising the Venus fly-trap domain (VFT) and the cysteine rich domain (CRD)) and the seven transmembrane domain (7TM). Orthosteric ligand binding sites are found in the cleft between the upper and lower lobes (black circles) of the VFT and the majority of allosteric modulators bind to sites in the 7TM (blue circles). Activation leads to a closure of the VFTs and a reorientation of the ECDs, the CRDs and the 7TMs bringing the two subunits into closer proximity. In N-terminally SNAP-tag labeled receptor dimers this leads to a transition from a high FRET/resting to a low FRET/active state. G protein activation through interactions with the cytoplasmic side of the 7TM is reported to occur at >10 ms timescales. The shown structures were obtained by homology modeling using mGlu5 structures PDB ID 6N52 and 6N51.

In the present study, we examine the effects of the 7TM-targeting mGlu2 PAM BINA on the conformational dynamics of the receptor at the single molecule level. Although a few studies reported on the structural dynamics of mGlu receptors on single molecules (Vafabakhsh et al, 2015; Levitz et al. 2016; Habrian et al. 2019; Liauw et al. 2021), none examined the allosteric modulation by small molecules or G proteins. Here, we optimized the conditions to conserve PAM activity at the solubilized full-length mGlu2 receptor and measure single molecule Förster resonance energy transfer (smFRET) at nanosecond time resolution. We show that mGlu2 is oscillating between inactive and active states at submillisecond timescales in its apo state, and that Glu partially increased the fraction of receptors residing in the active state. Only in the presence of Biphenyl-indanone A (BINA) can the full population of receptors be stabilized in an active conformation, providing a striking explanation for the increased agonist efficacy and potency observed with this PAM. We observe a similar effect with the nucleotide-free Gi heterotrimeric protein. Altogether, the quantification of submillisecond structural dynamics of soluble, functional, full-length mGlu2 receptors sheds new light on the mechanism of action of a synthetic mGlu2 PAM and the stabilizing effect of the Gi protein.

## Results

### Optimization of detergent conditions to obtain fully functional mGlu2 dimers

Our approach to perform smFRET measurements with submillisecond resolution requires fluorescently labeled receptors to be freely diffusing in solution, while maintaining full functional integrity for several hours at room temperature. Therefore, we evaluated a set of different detergents commonly used for GPCR-solubilization, supplemented or not with the cholesterol analogue cholesteryl hemisuccinate (CHS), for their ability to extract receptors from membranes and maintain them in solution, while preserving native-like ligand responsiveness. For this initial detergent screening we employed lanthanide resonance energy transfer (LRET) (Selvin 2002; Scholler et al. 2017), that monitors the ECD N-termini reorientation upon activation (**Fig. 1**), and was previously reported as an efficient approach to study mGlu structural dynamics (Doumazane et al. 2013; Olofsson et al. 2014; Scholler et al. 2017; Tora et al. 2018). We labeled the extracellular N-termini of the mGlu2 subunits using the SNAP-tag technology on HEK293T cells (**Fig. 2a**). This approach does not interfere with receptor function and by using cell-impermeable SNAP-tag substrates only cell surface receptors are labelled, resulting in a homogenous, fully processed, dimeric, fluorescently labeled receptor population (Doumazane et al. 2013).

**Figure 2:**
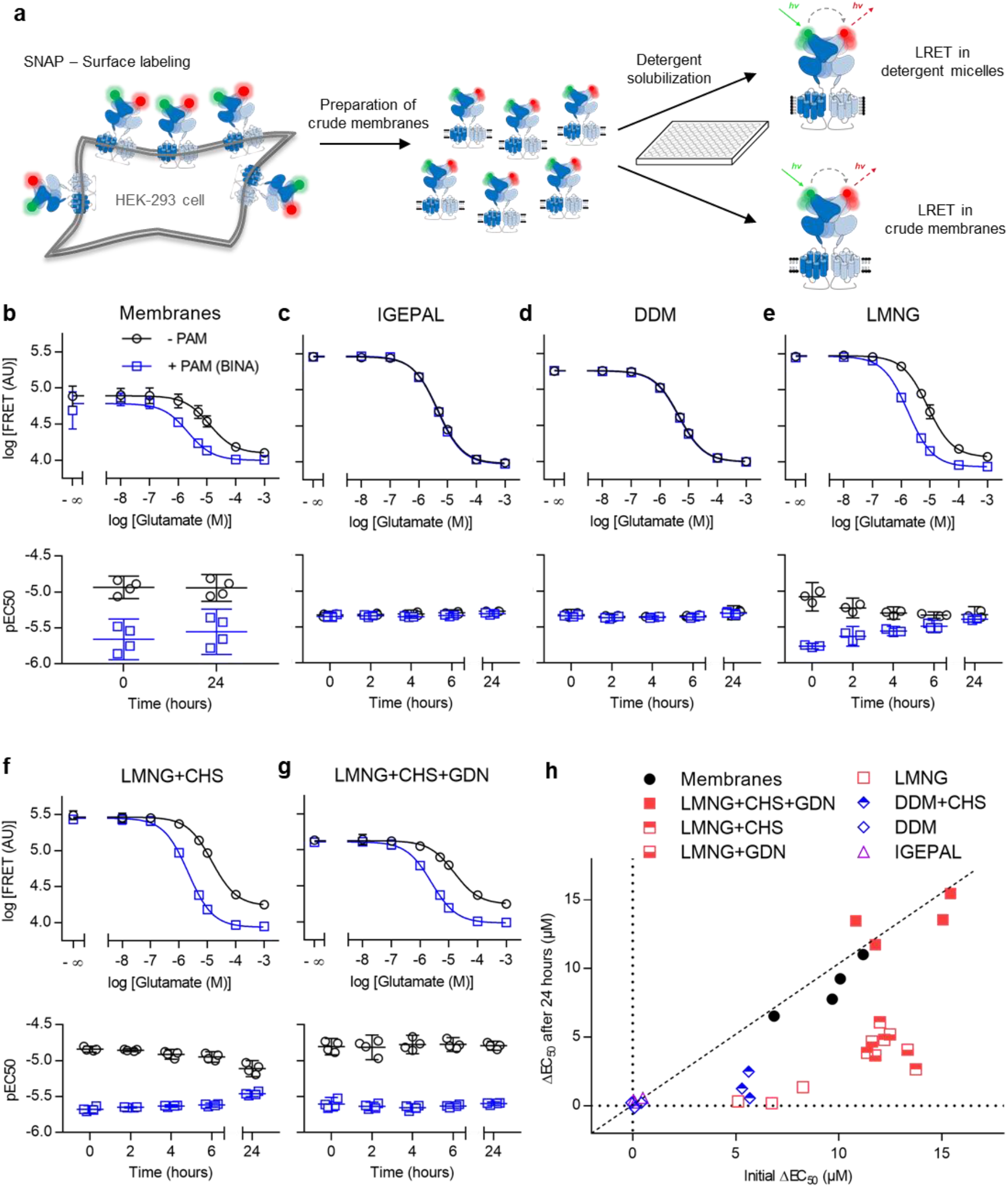
Evaluation of detergents for functional solubilization of full-length mGlu2 using LRET. **a)** SNAP-mGlu2 dimers were labeled with cell-impermeable lanthanide donor and green acceptor fluorophores on living HEK-293T cells. After preparation of crude membrane fractions, LRET measurements were performed in microtiter plates either directly on membranes or after detergent solubilization. **b-h)** The functional integrity of SNAP-labeled receptors was monitored over time at room temperature based on the dose-dependent intersubunit LRET changes in response to the orthosteric agonist Glu (- PAM) and in combination with 10μM positive allosteric modulator BINA (+ PAM). **b-g)** Dose-response curves at time 0h (top) and timecourse of pEC_50_ values (bottom) obtained on crudes membranes **(b)**, in IGEPAL **(c)**, DDM (**d**), LMNG (**e**), LMNG-CHS (**f**) and LMNG-CHS-GDN **(g)**. Data represent the mean of triplicate analysis from 3-4 biological replicates. Errors are given as standard deviation of the mean in dose-response curves **(b-g, top)** and the 95% confidence interval of the mean for pEC_50_ values **(b-g, bottom)**. **h)** Scatter plot of ΔEC_50_, the difference in EC_50_ obtained in presence and absence of BINA, at time 24h at RT (y-axis) vs. at time 0h (x-axis), for membrane fractions and detergent mixtures. The conditions along the diagonal represent those experiencing the lowest changes over time.

The functional integrity of receptor preparations was assessed upon Glu stimulation in detergent micelles, after various time points up to 24 h at room temperature, and compared with control conditions of mGlu2 in crude membranes. In parallel, the integrity of the transmembrane domain was evaluated through the effect of BINA (**Fig. 2b**). Indeed, the PAM-binding site is known to be located within the 7TM region (O’Brien et al. 2018) and thus the functional link translating the PAM effect to Glu potency at the ECD level provides a reliable measure of the receptor’s global functional integrity.

A dose-dependent response to Glu, reflected by a decreasing LRET signal, was observed under all tested conditions but in some cases revealed changes in the Glu pEC_50_ values over time (**Fig. 2b-h and S1-11**). More importantly, the effect of BINA on Glu potency strongly depended on the detergent mixture used, thus indicating a detergent-dependent integrity of the functional link between ECD and 7TM. Positive allosteric modulation was not observed using IGEPAL (octylphenoxypolyethoxyethanol) and DDM (n-dodecyl β-D-maltoside), two nonionic detergents that have previously been used to solubilize full-length mGluRs for smFRET by TIRF microscopy (Vafabakhsh et al. 2015) (**Fig. 2c-d, h, S1 and S2**). Only a weak allosteric modulation by BINA was found when DDM was supplemented with CHS, known to facilitate functional GPCR solubilization through mixed sterol-detergent micelles (Thompson et al. 2011), but this effect was lost within 4-6 hours (**Fig. 2h, S3**). Similarly, allosteric modulation in micelles composed of the branched nonionic detergent LMNG (lauryl maltose neopentyl glycol) was not stable (**Fig. 2e, h, S4**). In contrast, addition of CHS to LMNG led to a prolonged functional integrity of the receptors, lasting from 6 to 24 hours, in a CHS-dose-dependent fashion (**Fig. 2f, h, S5-7**).

The functional integrity of mGlu2 was further improved by the addition of GDN (glyco-diosgenin) to the LMNG-CHS mixture (**Fig. 2g, h, S8-10**). This steroid-based amphiphile has been demonstrated to improve GPCR stability (Chae et al. 2012) and was recently employed in structure determination of mGluR5 by cryo-EM (Koehl et al. 2019). GDN was found beneficial at all concentrations tested (**Fig. S8-10**), but the presence of CHS remained crucial for long-term functional integrity of solubilized receptors in micelles (**Fig. 2h, S11**).

Overall, our results demonstrated that the optimized LMNG-CHS-GDN mixture (0.005% w/v, 0.0004% w/v and 0.005% w/v, respectively) is mandatory to maintain the functional integrity and allosteric link between the mGlu2 ECD and 7TM domains, for at least 24 h at room temperature (**Fig. 2e, h, S9**). The LRET signal range as well as the pEC_50_ values for Glu and Glu + PAM in this mixture were well in agreement with those obtained in crude membranes (**Fig. 2b and S12**), and also reflected earlier observations in live cells (Doumazane et al. 2013; Olofsson et al. 2014). Interestingly, optimal GDN and CHS concentrations remained moderate, which turned out to be advantageous for our smFRET studies as both chemicals were slightly contaminated with fluorescent species of unknown origin (also found in batches from different suppliers).

### Allosteric modulation through the 7TM is required to stabilize the fully active ECD state

We then turned to the single molecule study of full-length mGlu2 and therefore substituted the LRET fluorophores with SNAP-tag substrates of Cy3B as donor and d2 as acceptor. Thanks to the PIE/nsALEX confocal configuration, which we previously employed to study isolated mGlu ECDs (Olofsson et al. 2014; Tora et al. 2018), single molecules are detected as they diffuse through the confocal observation volume **(Fig. 3a)**. Only donor-acceptor (D-A) containing complexes are selected based on the stoichiometry factor *S_PR_* (Hellenkamp et al. 2018; Kapanidis et al. 2004). For each single molecule, we further determined its apparent FRET efficiency (*E_PR_*), the average fluorescence lifetime of the donor in presence of the acceptor (*τ_DA_*) and the average excited state lifetime of the acceptor (*τ_A_*).

**Figure 3:**
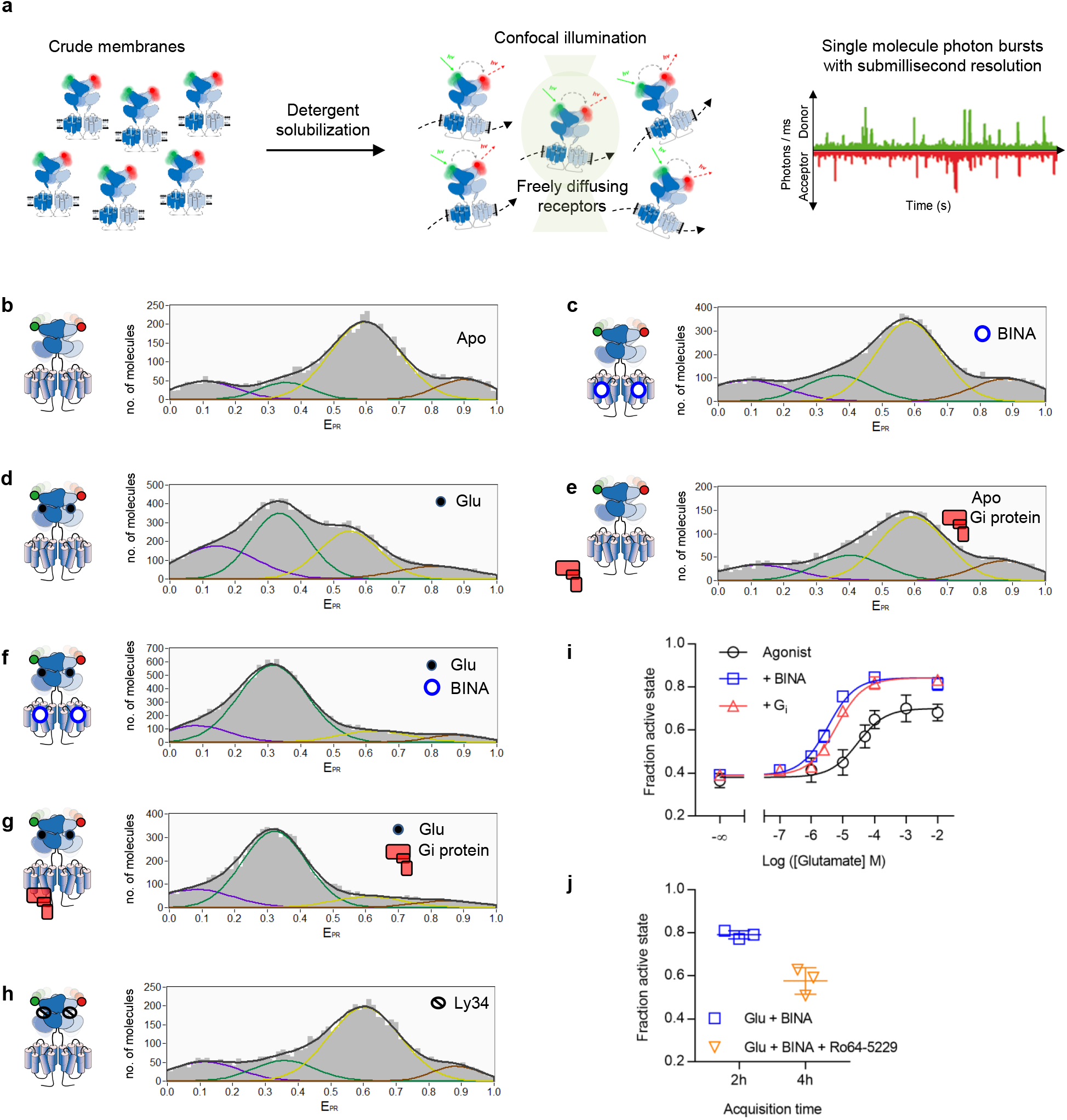
smFRET reveals the conformational landscape of full-length mGlu2 in LMNG-CHS-GDN micelles. **a)** SNAP-mGlu2 dimers were labeled with cell-impermeable Cy3b donor and d2 acceptor fluorophores on living HEK-293T cells. Then mGlu2 dimers were detergent-solubilized from crude membrane fractions and smFRET measurements were performed on freely diffusing molecules with confocal illumination. **b-h)** Representative histograms displaying the number of doubly labeled molecules as a function of apparent FRET efficiency (E_PR_). Distributions were obtained in the absence of ligand (Apo) or in the presence of Glu (10 mM), competitive antagonist LY341495 (1 mM), BINA (10 uM) and G protein (1 μM), as indicated. Colored lines represent Gaussian fitting, black lines correspond to the cumulative fitting (see text). All histograms revealed four major populations at very low FRET (VLF, purple), low FRET (LF, green), high FRET (HF, yellow) and very high FRET (VHF, red). **i)** smFRET analysis of the effect of Glu without (Agonist) or with BINA (10 μM) or G_i_ (1 μM), as indicated. The fraction of the active state is defined as the fraction of molecules in the LF+VLF populations over all molecules. **j)** smFRET analysis of the reversibility of the PAM-induced full ECD activation (500 nM, 2h) through competition with an excess of the NAM Ro64-5229 (10 μM, 4h). **(i-j)** show the data from three independent biological replicates with mean and standard deviation.

FRET histograms of SNAP-labeled, full-length mGlu2 in LMNG-CHS-GDN micelles showed a wide, multimodal distribution (**Fig. 3b-h**), indicating the co-existence of four main ECD states. In the absence of ligands, a main population was centered around *E_PR_* ~ 0.6 (high FRET, HF, yellow), and less well-defined minor populations were present at lower and higher FRET values (**Fig. 3b**). Upon application of saturating concentrations of Glu, a second major population at low FRET (LF, *E_PR_* ~ 0.34, green) appeared (**Fig. 3d**). Such a decrease in FRET was observed in our smFRET study on freely diffusing isolated VFTs (Doumazane et al. 2013; Olofsson et al. 2014), as well as on immobilized full-length receptors (Vafabakhsh et al. 2015). Nevertheless, in contrast to our observation on isolated VFTs, which showed a complete shift of the major population to lower FRET, a portion of the HF population remained for the full-length receptor.

Next, we explored the effect of a PAM at saturating concentrations. Alone, BINA had no effect (**Fig. 3c**), which agrees with the expected effect of a pure allosteric modulator that requires an agonist to reveal its modulatory activity. Thus, BINA does not act as an ago-PAM with regard to ECD reorientation. In contrast, in presence of saturating Glu, BINA unveiled its PAM effect, and led to a strong increase of the LF population (**Fig. 3f**) accompanied by a nearly complete depopulation of the HF states. We then analyzed the influence of the heterotrimeric G_i_ protein, known to stabilize For mGlu2, it is established that the G_i_ protein influences the ECD reorientation (Doumazane et al. 2013). Interestingly, presence of the nucleotide-free heterotrimeric G_i_ complex at saturating concentrations led to nearly identical FRET distributions as those promoted by BINA, both in absence and presence of Glu (**Fig. 3e and g**, respectively). The combination of BINA and G_i_ did not result in a further detectable synergistic effect (**Fig. S15a**). We therefore concluded that BINA as well as G_i_ exert an allosteric control through the 7TM, which is required for a complete reorientation of the ECD toward the LF state.

Finally, we noted that application of saturating concentrations of the competitive orthosteric antagonist LY341495 (LY34) led to a similar distribution as seen for the apo receptor (**Fig. 3h**). Thus, no basal receptor activity or residual Glu was observed in our preparations, which was further verified by titration with LY34 in LRET measurements (**Fig. S13a-e**).

The major changes in response to ligands were found to result from depopulation of the HF state accompanied by an increase of the LF state, while no notable changes were observed in the two minor populations found at very low FRET (VLF, *E_PR_* ~ 0.1, purple, see also **Fig. S14d**) and very high FRET (VHF, *E_PR_* ~ 0.87, red, see also **Fig S14c**). We assigned the VHF and HF populations to a conformational ensemble representing the resting/inactive VFT state and the VLF and LF populations to the active state (**Fig. 3b-h**). Then, to gain a quantitative view of mGlu2 receptor activation, we calculated the fraction of active molecules found in the LF+VLF states relative to all molecules. We fitted all distributions with four Gaussians, and recovered similar values of *E_PR_* and full width half maximum (FWHM), pointing to the fact that similar FRET states are populated for all conditions tested (**Fig. S14a-b**). The fractional amplitude of the LF+VLF states recovered from the fit allowed us to plot dose-response curves. pEC_50_ values obtained for Glu in the absence or the presence of saturating concentrations of BINA (**Fig. 3i**) were in good agreement with those obtained from ensemble LRET on membranes (**Fig. 2b**) or in optimized detergent micelles (**Fig. 2g**). The allosteric effect of BINA on the apparent Glu potency (an increase by almost one order of magnitude) as well as its effect on the maximum efficacy were also recovered (**Fig. 3i**). This effect was reversible, as addition of an excess of the negative allosteric modulator (NAM) Ro64-5229 (Ro64) to receptors after activation by 500 nM BINA + Glu decreased the fraction of active receptor to a similar level observed in the presence of Glu alone (**Fig. 3j**). Altogether, these results further validated the full functional integrity and native-like ligand responsiveness of our receptor preparations in optimized detergent micelles.

In addition, Glu titration at saturating G_i_ concentration was strikingly similar to the one obtained with BINA (**Fig. 3i**). Thus, G_i_ acts as an allosteric modulator on Glu potency and ECD activation. Most notably, no additional populations or substantial changes in the four major peak positions *(E_PR_)* were found in the presence of BINA or Gi. This indicates that even if BINA and G_i_ promote alternative conformations through distinct interaction sites at the 7TM level, their allosteric effect on the ECD dimer conformation can be explained by a simple shift of the equilibrium toward the active state, rather than the stabilization of alternative states.

### BINA or G_i_ are required to suppress submillisecond dynamics and stabilize the active state

We then took advantage of the high time resolution of our PIE/nsALEX approach to uncover hidden states, sampled by the receptor during its residence time in the confocal illumination volume (here ~5 ms). Interconversions between multiple FRET efficiency states at timescale faster than this residence time lead to averaging, which results in populations being found at intermediate FRET efficiency values when calculated by integrating over the entire residence time.

We employed two different methods to gain insights into the dynamic behavior of the mGlu2 ECD in full-length receptors. First, we plotted donor fluorescence lifetimes τ_DA_ for each single molecule against the γ-corrected FRET efficiency *E* (“τ_DA_ vs. *E*” analysis (Sisamakis et al. 2010)). This representation allows to identify structural dynamics, if populations deviate from the theoretical “static FRET line” (yellow line, **Fig. 4a-c**). For apo receptors, the main HF population appeared above the static FRET line **(Fig. 4a)**, thus indicating submillisecond conformational oscillations. In contrast, the LF population promoted by application of Glu (**Fig. 4b**) and further populated in the presence of Glu + BINA (**Fig. 4c**) was found much closer to the static FRET line, therefore implying reduced dynamics of the active ECD state.

**Figure 4:**
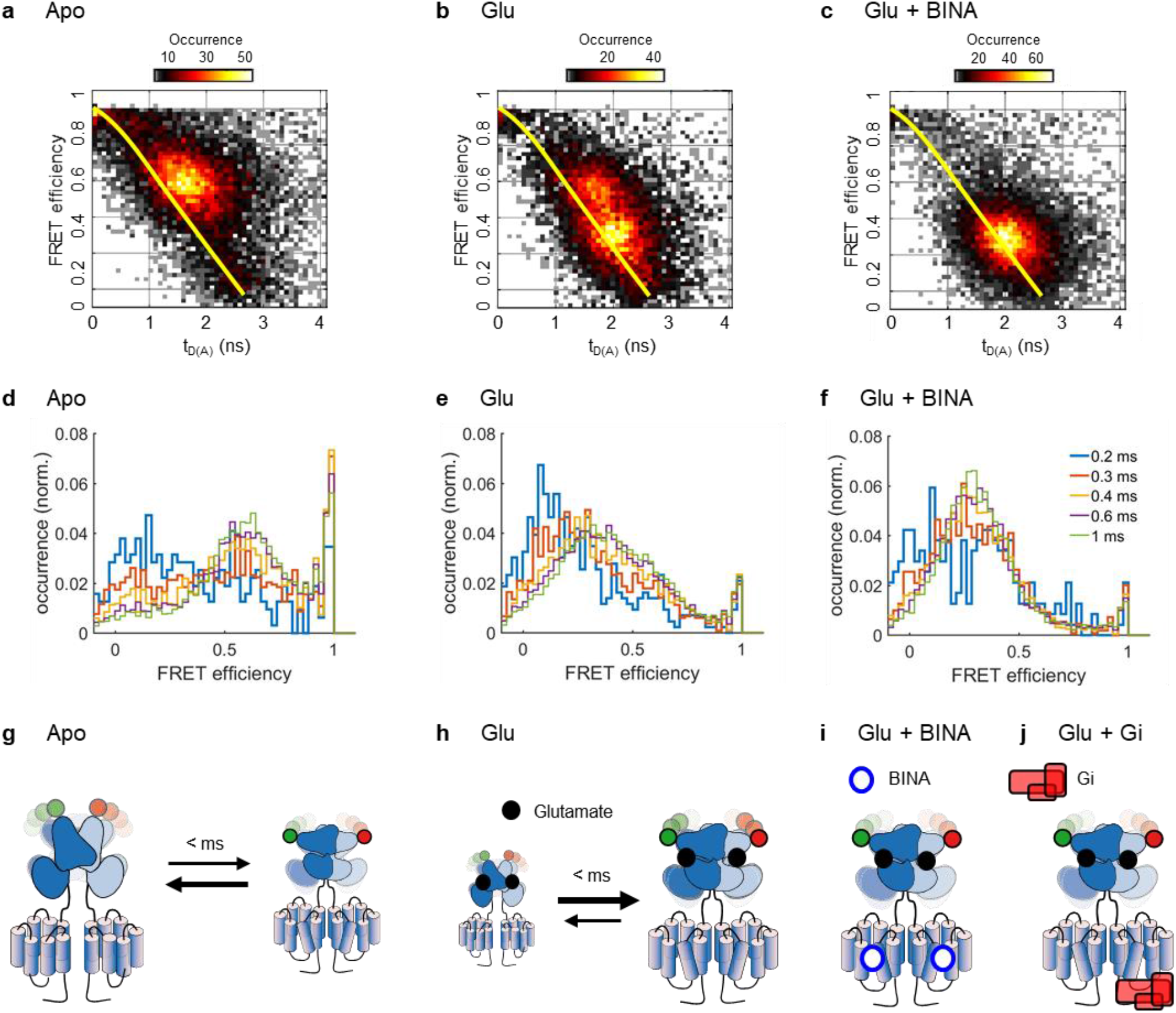
Structural dynamics analysis of mGlu2 dimers in response to orthosteric and allosteric ligands. **a-c)** Representative tDA vs. *E* histogram for mGlu2 dimers in the absence **(a)** or presence of Glu **(b)** or Glu + BINA **(c)**. For the Apo receptors, the major population deviates from the “Static FRET” line (yellow), indicating conformational dynamics at the sub-millisecond time scale. Addition of Glu stabilizes the receptor ECD in an ensemble of low FRET conformations with less flexibility, an effect that is reinforced by the allosteric modulator BINA. **d-f)** Time windows analysis for different integration times (from 0.2 to 1ms) reveals a large conformational flexibility of the Apo receptor ECD at 200-600μs timescales, that is strongly restricted when bound to orthosteric agonist and allosteric modulator**. g-j)** Schematic representation of the major species observed with different ligands, with the timescales of the transition between them **(g:** Apo**, h:** Glu**, i:** Glu+BINA**, j:** Glu + Gi). The size of the cartoons schematically represents the fraction of the species under the given condition.

Second, we performed time windows analysis (TWA) (Gopich and Szabo 2010), which relies on recalculating the FRET efficiency at integration times shorter than the residence time, here in 200 μs steps from 1 ms down to 200 μs. Shortening the integration time below 300 μs strikingly led to the disappearance of the main HF population for apo receptors, while two populations at *E* ~ 0.2 and >0.9 were revealed (**Fig. 4d**, red). This indicates that at integration times longer than 300 μs the apparent FRET population centered at *E* ~ 0.6 represents the time-averaged FRET value between these two sampled states. We therefore conclude that at submillisecond timescales, the apo receptor samples a set of conformations at low and very high FRET values, representing the active and inactive states, respectively (**Fig. 4g**). Of note, the distribution obtained by integration at 1 ms (**Fig. 4d**, green) matched the one obtained from calculations integrated over the entire residence time (**Fig. 3b**), indicating no detectable dynamics between 1 ms and the residence time of ~5 ms. Addition of the antagonist LY34 led to a distribution similar to the apo state (**Fig. S17-18**), thus excluding a stabilization of the oscillating ECDs by this orthosteric ligand.

In contrast, orthosteric and allosteric agonists promoted a stabilization of the ECDs in an ensemble of low FRET conformations (**Fig. 4e-f**). This was particularly obvious in the presence of Glu + BINA, where the majority of molecules remained within the LF population at *E*~0.25 even at lower integration times. Indeed, only some residual conformational dynamics, limited to low FRET states, were observed at an integration time of 200 μs (**Fig. 4f**, blue). A similar stabilization was observed for Glu-activated receptors in the presence of Gi, underlining the close similarity of allosteric modulation exerted by PAM and G protein on ECD dynamics (**Fig. S17-18**). Altogether we concluded that the synergistic binding of Glu and an allosteric modulator, either the G protein or a small molecular synthetic PAM, promoted the stabilization of the mGlu2 receptor in an ensemble of conformations characteristic of the active state, stable for at least several milliseconds as given by the residence time in the confocal volume (**Fig. 4i-j**). In the case of receptors bound to Glu alone, this stabilizing effect of the active state was less pronounced (**Fig. 4b and e**), with a fraction of receptors still sampling between the high FRET resting state and the low FRET active state (**Fig. 4h**). This confirms the inability of the natural full agonist Glu to fully stabilize the ECDs in the active state and emphasizes the importance of a long-range functional link between the 7TMs and the ECDs that allows for allosteric interdomain communications mandatory for maximal stabilization of the ECDs in the active state.

### Loss of functional integrity and dynamics of mGlu2 in IGEPAL

Our LRET data demonstrated that receptors only provide a strong and stable response to PAM in LMNG-CHS-GDN micelles, while in IGEPAL or DDM micelles, even supplemented with CHS, this effect was completely absent (**Fig. 2**, **S1**). To further understand the differential effects of these detergent mixture, we analyzed the effect of Glu, BINA, G_i_ and Ro64 on receptors solubilized with IGEPAL by smFRET (**Fig. S16**). In contrast to the data obtained in LMNG-CHS-GDN, we found that Glu was sufficient to totally stabilize the receptors in the active ECD state (**Fig. S16**, see also dynamics analysis, **Fig. S17-S18**), similarly to earlier reports(Vafabakhsh, Levitz, and Isacoff 2015). No further effect on ECD activation was observed upon addition of BINA or the G_i_ protein and likewise the NAM Ro64 was not capable of reducing the fraction of the active state (**Fig. S16**) in a way seen in LMNG-CHS-GDN. These observations underline the loss of allosteric effects in IGEPAL or other detergents mixtures. Such lack of functionality of the receptor could arise from: 1/ a loss of allosteric communication between the 7TM and the ECD; 2/ a loss of structural integrity of the 7TM that becomes unable to bind the PAM, NAM and G protein, or 3/ a direct effect of IGEPAL on the conformation of the 7TM, stabilizing it in a PAM-bound-like conformation, that should only be reached in the presence of the allosteric modulator under native-like conditions.

### Maximal ECD activation remains ligand-dependent

Next, we addressed the mode of action of partial agonists, previously shown to promote changes in the ECD intersubunit orientation but to a lower extent than full agonists (Doumazane et al. 2013). Pharmacologically, partial agonists are ligands that do not trigger maximal cellular responses, not even at saturating concentrations (Rosenbaum, Rasmussen, and Kobilka 2009). At the structural level, this may either be explained by the existence of specific intermediate active conformations (Masureel et al. 2018) or by a less efficient shift of the resting-to-active equilibrium compared to full agonists. Our previous data proposed a simple shift in the equilibrium of isolated ECD dimers rapidly oscillating between active and resting conformations toward the active state, while maintaining submillisecond dynamics (Olofsson et al. 2014). Here, in full-length receptors in LMNG-CHS-GDN, the *E_PR_* peak positions of the four populations described in Figure 3 were perfectly recovered for the partial agonists LCCG-I (**Fig. 5a**), DCG-IV (**Fig. 5b**) and LY354740 (LY35) (**Fig. S14-15**). Nevertheless, the extent of depopulation of the HF state and corresponding increase in the population of the LF state remained ligand-dependent. Quantification of the fraction of activation indicated that these molecules have a lower efficacy than Glu to populate the active state (**Fig. 5h-i**). We further observed submillisecond dynamics of the HF state under these conditions, pointing to the inability of these partial agonists to efficiently stabilize the less dynamic active ECD state. Indeed, the HF population appeared above the static FRET line, while the FRET distributions in TW analysis remained intermediate between those of the apo and the Glu-bound receptors (**Fig. S19**). Addition of BINA (**Fig. 5d-e**) or G_i_ (**Fig. 5f-g**) further pushed the equilibrium toward the active state, but to a lower extent than obtained with Glu (**Fig. 5h-i**). This observation revealed that these partial agonists are unable to fully stabilize the ECD in the active orientation, even in the presence of BINA or the heterotrimeric G_i_ and consequently, that maximal ECD activation still remains dependent on the individual efficacy of an agonist. Furthermore, these results together with the finding that all studied conditions resulted in the same four major FRET states (**Fig. S14a**) demonstrate that partial agonists do not stabilize intermediate FRET states but shift the equilibrium between the dynamic inactive and the less dynamic active ECD states.

**Figure 5:**
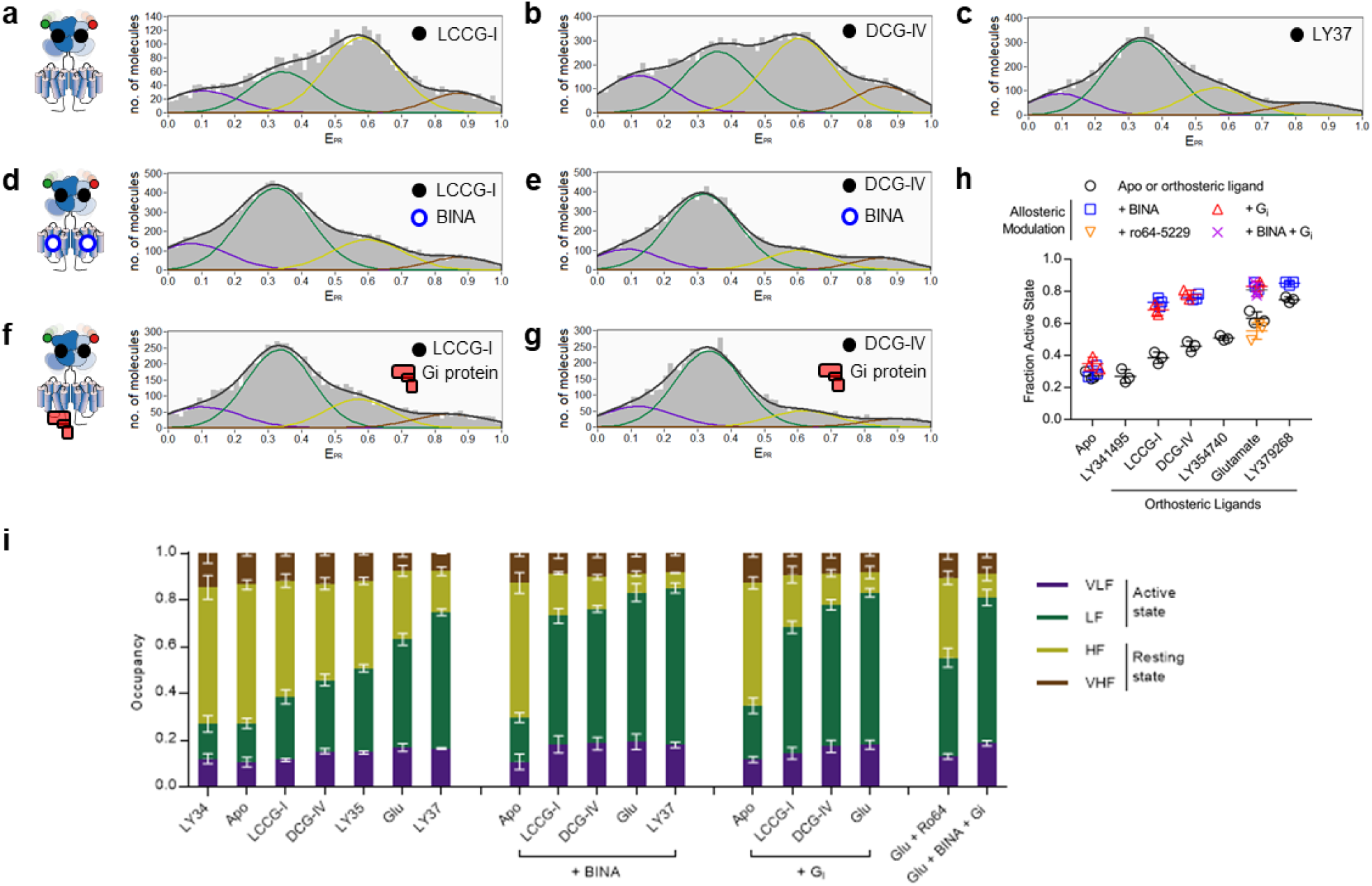
Different efficacies of orthosteric ligands on mGlu2 ECD rearrangement. **a-g)** FRET distributions were obtained in the presence of partial agonists LCCG-I **(a)**, DCG-IV **(b)**, synthetic full agonist LY379268 **(c),** LCCG-I + BINA **(d)**, DCG-IV + BINA **(e)**, LCCG-I + Gi **(f)** and DCG-IV + Gi **(g)**. **h)** Comparison of the fraction of the active state in response to different orthosteric and allosteric ligands. The scatter plot shows the data from three independent biological replicates with mean and standard deviation. **i)** Comparison of the fraction of all states in response to different orthosteric and allosteric ligands. The error bars represent the standard deviation from three independent biological replicates.

### The natural full agonist glutamate does not exhibit maximal efficacy

Finally, we further characterized the synthetic full agonist LY379268 (LY37) at the single molecule level. Interestingly, this ligand appeared more potent than Glu to stabilize the ECD in its active state (**Fig. 5c, h, i**). The E_PR_ histogram showed a higher fraction of molecules in the active state than for Glu. Similarly, the dynamics analysis revealed a stabilization of the majority of molecules in the LF states, for up to several milliseconds (**Fig. S17-S18**). This observation points to the possibility that LY37 might qualify as a superagonist (Schrage et al. 2016), i.e. a compound that displays greater efficacy and thus higher receptor signaling output, than the endogenous full agonist Glu. However, this effect is only observed when the receptor is solely bound by the orthosteric ligand, as the distribution of states obtained upon activation in the presence of PAM was identical for receptors bound by Glu and LY37 (**Fig. 5h-i and S15**).

## Discussion

GPCR activation can be finely tuned by different classes of ligands acting either via orthosteric or allosteric sites. Among them, PAMs enhance agonist action by increasing their potency and/or efficacy. Here, we used smFRET to explore how a PAM can increase the efficacy of mGlu2 receptors, by monitoring the fast dynamics of the intersubunit rearrangement of the ECDs. We analyzed the effect of BINA, an mGlu2 specific PAM, on isolated, full-length and fully functional receptors with submillisecond time resolution, relevant for the conformational movements of such protein domains (Henzler-Wildman and Kern 2007). Our data reveal the presence of four ECD states, two of which - the HF/inactive and LF/active states - are predominantly populated in a ligand-dependent manner. Two minor populations (VHF and VLF) were barely affected by ligands, but we note that they could be in exchange with the two major populations at timescales slower than the resolution of our method (> 5 ms).

The conformational landscape of receptor populations clearly differed from the one observed with the isolated ECD dimers (Olofsson et al. 2014). In that latter case, all dimers were shown to be oscillating at a ~100 μs timescale between high and low FRET states, in response to all ligands tested. Here, in the case of full-length receptor dimers in the apo state or bound to antagonist, the main population is similarly oscillating between the HF and LF states (at a slightly slower timescale of ~200-500 μs). However, in contrast to the isolated VFT dimer, addition of full agonist led to a stabilization of an ensemble of LF/active states, an effect further promoted by a PAM. These states appear stable for at least several milliseconds, a duration compatible with the activation of downstream signaling (Marcaggi et al. 2009; Grushevskyi et al. 2019). We propose that this stabilization of the active ECD state stems from a strengthening of the active dimeric interface, probably via interactions involving transmembrane helix 6, as reported based on crosslinking experiments at the surface of live cells (Xue et al. 2015; 2019) and structure determination for mGlu5 (Koehl et al. 2019), mGlu1(Zhang et al. 2020) and GABA_B_ receptors (Shaye et al. 2020).

Saturating concentrations of partial agonists or of the natural agonist Glu were not able to fully depopulate the basal HF state and stabilize receptors in the active LF state during the observation time of several milliseconds. Addition of BINA to the partial agonists was not sufficient to promote the stabilization of the active state to the extent observed with the full agonists Glu and LY37. Thus, the extent of activation remains ligand-dependent even in the presence of allosteric modulators. In contrast, LY37, formally considered a full agonist like Glu, appears more efficient than Glu in promoting the active ECD state in our assay, which qualifies this molecule as a “superagonist” (Schrage et al. 2016). Likely, this effect was previously hidden in cell-based assays, due to amplification of the signaling cascade and saturation of the readout (Doumazane et al, 2013).

The presence of the nucleotide-free heterotrimeric G_i_ protein complex was found to produce the same effect as the PAM, allowing Glu to fully stabilize the active state. Like BINA, G_i_ also increased partial agonist efficacy in populating the active state. Of note, the effects of BINA and G protein are not additive, suggesting they exert a similar effect. This is consistent with our observation that the G protein-bound state of another class C GPCR (the GABAB receptor), is similar to that observed with an agonist and a PAM (Shaye et al. 2020, and our unpublished observation). Our data are also consistent with the positive allosteric action of G proteins on GPCRs (Xiao et al. 2009), as also observed with class C receptors, including mGlu2 (Doumazane et al. 2013). Within the cellular environment, such an effect is expected to be transient, as upon GTP binding, the receptor-G protein complex dissociates and the allosteric action is lost. As such, PAMs can maintain this effect by stabilizing receptors in their active G protein bound-like state, which then facilitates G protein binding and activation.

Our data contradict those previously reported, showing an apparent full stabilization of the mGlu2 receptor in its active state with Glu alone in IGEPAL conditions (Vafabakhsh et al. 2015; Levitz et al. 2016; Habrian et al. 2019). It is possible that the lower time resolution in these assays prevents the detection of the basal state population. However, our results obtained in these conditions of detergents suggest that the 7TM domains behaves as if they were already occupied by a PAM, likely already being in an active-like state. Interestingly, a recent study using smFRET on a sensor reporting on the proximity between the CRD domains suggested that the fully active conformation of the receptor could not be reached in the presence of Glu alone (Liauw et al. 2021). A stabilization in the fully active state required the C770A mutation in the 7TM domain, described to enhance mGluR signaling in a manner similar to a PAM. Although this observation supports our model that a PAM effect is required for the full stabilization of the mGluR2 active state, we note that the effect of the binding of a synthetic PAM or of the G_i_ protein was not described in that study. It is likely that the receptor was in fact not able to be activated by such PAM under the detergent conditions used (DDM (0.05%) + CHS (0.005%)), for which we report here the absence of effect of BINA under very similar detergent conditions (**Fig. S3**).

Therefore, our results and the comparison with previous studies demonstrate that a careful optimization of solubilization conditions is required to maintain the functional integrity of full-length mGlu2 at room temperature. This was only achieved using a mixture of LMNG-CHS-GDN, while all other tested conditions employing popular detergents exhibited a time-dependent impact on receptor function. It is not surprising that a functional reconstitution of the multidomain, multimeric mGlu receptor requires adapted characteristics to account for proper folding, ligand binding and activity. Improved functionality by the branched nonionic detergent LMNG through enhanced stabilization of the 7TM (Lee et al. 2020) and beneficial polar interactions of the maltoside head with loops and 7TM ends may play an important role in maintaining the functional link between the ECD and the 7TM (Koehl et al. 2019). Further stabilization and functionality are provided by the two sterol-containing compounds CHS and GDN. Both mimic cholesterol, known to be important for class C GPCR function (Wu et al. 2014; Huang et al. 2016) and CHS further provides a net negative charge to the detergent micelles. Negative charges have been described to enhance agonist affinity and stabilize the active state of the β2-AR (Strohman et al. 2019), a prototypical class A GPCR, whose orthosteric binding site comprises similar features to that of the allosteric site in mGlu (Feng et al. 2015). Nevertheless, the triple combination LMNG-GDN-CHS was required to maintain receptor function over time, pointing to a complementary role of these molecules. Taken together, these observations highlight the importance of the lipid environment on mGlu receptor function.

Overall, by identifying conditions under which the solubilized mGlu2 receptor conserves its modulation by a PAM and the G protein, we have been able to show that BINA can increase the population of active receptors in the presence of Glu. The increased efficacy observed with this PAM arises from its ability to stabilize the active state already populated in the presence of orthosteric agonists. Our study paves the way for a deeper understanding of how the structural dynamics of mGlu receptors as well as other membrane receptors regulate their function and may open up new routes for the development of fine-tuned therapeutics.

## Acknowledgements

We thank Sebastien Granier and Rémy Sounier (IGF, Montpellier, France) for providing the heterotrimeric G protein; Gilles Labesse (CBS, Montpellier) for helping in mGlu2 modelling; Suren Felekyan (U. Dusseldorf, Germany) for discussion on data analysis; Guillaume Lebon, Alexandre Bouyssou (IGF, Montpellier, France) and the members of the IBM team (CBS, Montpellier) for fruitful discussions; the Arpège platform (IGF, Montpellier) for providing facilities and technical support, Perkin Elmer Cisbio for providing reagents. Our research is supported by grants from the Agence Nationale pour la Recherche (ANR 17-CE09-0026-02 to EM and ANR 18-CE11-0004-02 to EM and JPP), and the Fondation Recherche Médicale (DEQ20170336747 to JPP). AMC was supported by the Labex EPIGENMED. The CBS belongs to the France-BioImaging national infrastructure supported by the French National Research Agency (ANR-10-INBS-04, “Investments for the future”) and is supported by the GIS “IBiSA: Infrastructures en Biologie Santé et Agronomie”.

## Authors contributions

Anne-Marinette Cao: Detergent screening, protein preparation and purification, preliminary LRET and smFRET experiments, revised the manuscript

Robert B. Quast: Detergent optimization, protein preparation and purification, LRET and smFRET data acquisition and analysis, wrote the manuscript

Fataneh Fatemi: Preliminary protein preparation and purification and smFRET experiments, revised the manuscript

Philippe Rondard: Experiment design, data interpretation, wrote the manuscript.

Jean-Philippe Pin: Experiment design, data interpretation, acquisition of funds, wrote the manuscript

Emmanuel Margeat: Experimental design, data analysis and interpretation, design and implementation of the smFRET setup, acquisition of funds, wrote the manuscript

## Materials and Methods

### Chemicals

All chemicals were purchased from Sigma-Aldrich, Merck and Roth unless otherwise noted. n-dodecyl-β-D-maltopyranoside (DDM), lauryl maltose neopentyl glycol (LMNG) and cholesteryl hemisuccinate (CHS) tris salt were purchased from Anatrace (through CliniSciences, France). Glyco-diosgenin (GDN) was purchased from Avanti Polar Lipids through Merck. SNAP-Lumi4-Tb, SNAP-green, SNAP-Cy3b and SNAP-d2 were obtained from Cisbio Bioassays (Codolet, France). DCG-IV, LY341495, LY379268, LY354740, LCCG-I, BINA hydrochloride and Ro64-5229 were purchased from Tocris Bioscience (Bristol, UK).

### Plasmids

The pcDNA plasmid encoding SNAP-tagged human mGluR2 was a gift from Cisbio Bioassays (Codolet, France).

### Cell culture

Adherent HEK293T cells (ATCC CRL-3216, LGC Standards S.a.r.l., France) were cultured in Gibco_™_ DMEM, high glucose, GlutaMAX_™_ Supplement, pyruvate (Thermo Fisher Scientific, France) supplemented with 10% (vol/vol) FBS (Sigma-Aldrich, France) at 37°C, 5% CO_2_ and passaged twice per week.

### Transfection and labeling

1×10^7^ cells were seeded in 75cm^2^ flasks 24 h prior to transfection with Polyethylenimine (PEI 25K, Polysciences Europe GmbH, Germany) at a DNA to PEI ratio (w/w) of 3:1 using 12 μg mGluR2 plasmid DNA per flask. In brief, 10 mg/ml PEI stock solution in 1 M HCl was diluted in 20 mM MES at pH 5 with 150 mM NaCl together with the DNA and incubated at room temperature for 25min before sequential addition of 5 ml complete medium followed an additional 5 ml. The flask’s culture medium was then replaced by the diluted transfection mix and protein expression proceeded for 48h at 37°C, 5% CO_2_.

SNAP-tag labeling was performed on surface-adhered cells in DMEM GlutaMax without FBS for 1-2h at 37°C using final concentrations of either 100 nM SNAP-Lumi4-Tb and 60 nM SNAP-green for LRET or 600 nM SNAP-Cy3b and 300 nM SNAP-d2 for smFRET measurements. Following labeling, excess dye was removed by three cycles of washing with DPBS w/o Ca^2+^ and Mg^2+^ (Thermo Fischer Scientific, France) at ambient temperature.

### Preparation of crude membrane fractions

Adherent cells were detached mechanically using a cell scraper in DPBS w/o Ca^2+^ and Mg^2+^ (Thermo Fisher Scientific, France) and collected at 500 x g and 22°C. Subsequently, cells were resuspended in cold hypotonic lysis buffer (10mM HEPES pH7.4, cOmplete_™_ protease inhibitor), frozen and stored at −80°C. After thawing, cells were passed through a 0.4mm gauge needle 30-times using syringe on ice. After two rounds of centrifugation at 500 x g and 4°C for 5min, the supernatant was aliquoted and centrifuged at 21,000 x g and 4°C for 30 min to collect crude membranes. The pellets were washed once with 20 mM HEPES pH 7.4, 118 mM NaCl, flash frozen in liquid N2 and stored at −80°C.

### Detergent solubilization

Receptors were solubilized on ice by resuspension of crude membranes in acquisition buffer (20 mM Tris-HCl pH7.4, 118 mM NaCl, 1.2 mM KH_2_PO_4_, 1.2 mM MgSO_4_, 4.7 mM KCl, 1.8 mM CaCl_2_) supplemented with 1% (v/v) IGEPAL, 1% (w/v) DDM, 1% (w/v) DDM + 0.2% (w/v) CHS Tris, 0.1% (w/v) LMNG, 0.1% (w/v) LMNG + 0.1% GDN (w/v), 0.1% (w/v) LMNG + 0.004%, 0.008% or 0.016% CHS Tris (w/v) or 0.1% (w/v) LMNG + 0.008% (w/v) CHS Tris + 0.05%, 0.1% or 0.2% GDN (w/v). After 5min, the solution was centrifuged for 10 min at 21,000 x g and 4°C. The supernatant was then applied to a Zeba Spin Desalting Column (7 kDa cut-off, Thermo Fisher Scientific, France) equilibrated in acquisition buffer containing 5% of the detergent concentration used for solubilization and centrifuged 2 min at 1,500 x g and 4°C. The flow-through was then immediately diluted 1:20 in cold acquisition buffer and kept on ice in the dark until use.

### LRET

Intersubunit LRET measurements of mGluR2 dimers, N-terminally labeled with the Lumi4-Tb donor and the green acceptor via an engineered SNAP-tag, were performed on a PHERAstar FS microplate reader (BMG Labtech, Germany) in white 384 well plates (polystyrene, flat-bottom, small volume, medium-binding, Greiner Bio-One SAS, France). Measurements where performed in acquisition buffer in the presence of indicated ligands at room temperature and plates where sealed and stored in the dark in between measurements for time course experiments to minimize evaporation and fluorophore bleaching. The fluorescence decay of donor and acceptor was recoded using the LRET 337/620/520 optical module by excitation with 20 flashes per well every 5 μs for a total of 2500 μs. The FRET signal was expressed as sensitized acceptor emission integrated between 50-100 μs and normalized to its emission between 1200-1600 μs as previously optimized for the given mGluR2 FRET sensor (Scholler et al. 2017).

### Expression and purification of heterotrimeric G_i1_

The heterotrimeric G_i1_ complex was a kind gift from Sebastien Granier and Remy Sounier (IGF Montpellier, France). G_i1_ heterotrimer was expressed in *Sf9* insect cells in EX-CELL 420 media (Sigma). Human G_αi1_ was cloned into the pVL1392 vector, and the virus was prepared using the BestBac system (Expression System, LLC). N-terminal Flag-tagged human G_β1_, and human G_γ2_ were cloned into the pFastBac vector, and the virus was prepared using the Bac-to-Bac baculovirus system. The cells were infected with both G_αi1_ and Gβγ virus at a ratio determined by small-scale titration experiments at 27°C for 48 h before collection. Cells containing G_i1_ heterotrimer were lysed in hypotonic buffer containing 10 mM Tris pH 7.4, 100 mM MgCl_2_, 10 mM GDP, 5 mM β-mercaptoethanol, and protease inhibitors. After centrifugation, membranes were dounced and solubilized in buffer containing 20 mM HEPES pH7.5, 100 mM NaCl, 1% DDM, 5 mM MgCl_2_, 10 mM GDP, 5 mM β-mercaptoethanol and protease inhibitors. Solution containing the G_i1_ heterotrimeric complex was loaded onto an anti-FLAG M1 affinity column. After washing of the column with 5 column volumes of buffer E1 (20 mM HEPES pH 7.5, 100 mM NaCl, 1% DDM, 5 mM MgCl_2_, 10 mM GDP, 5 mM β-mercaptoethanol) and buffer E2 (20 mM HEPES pH 7.5, 50 mM NaCl, 0.1% DDM, 1 mM MgCl_2_, 10 mM GDP, 100 μM TCEP) at a flow rate of 2 ml.min^-1^. After a detergent exchange was performed by washing the column with a series of seven buffers (3 CV each) made up of the following ratios (v/v) of MNG buffer (20 mM HEPES pH 7.5, 50 mM NaCl, 0.5% MNG, 1 mM MgCl_2_, 10 mM GDP, 100 μM TCEP) and E2 buffer: 0:1, 1:1, 4:1, 9:1, 19:1, 99:1 and MNG buffer alone. G_i1_ heterotrimer was eluted with Elution buffer (20 mM HEPES pH 7.5, 50 mM NaCl, 0.01% MNG, 1 mM MgCl_2_, 10 mM GDP, 100 μM TCEP). Eluted sample was concentrated in a 50 kDa MWCO concentrator to 100 μM and aliquots were flash frozen in liquid Nitrogen and stored at −80°C.

### PIE-MFD smFRET setup

Single-molecule FRET experiments with pulsed interleaved excitation (PIE) – multiparameter fluorescence detection (MFD) were performed on a homebuilt confocal microscope as described previously (Olofsson and Margeat 2013).

In brief, the 20MHz-pulsed white excitation laser was split into two beams spectrally filtered using excitation bandpass filters at wavelength 532/10 (prompt beam) and 635/10 (delayed beam) to excite the Cy3b donor and d2 acceptor molecules, respectively. The delayed beam has a path length of ~ 8m relative to the prompt beam, generating a ~24 ns delay in the pulse. The two beam paths are then recombined and focused using a 10x objective into a single-mode fiber, by which the beams become spatially overlapped and filtered. The output of the fiber is collimated using a 10 x microscope objective lens, polarized and coupled into an inverted microscope (Eclipse Ti, Nikon, France). The excitation power was controlled to give 25 μW for the prompt and 12 μW for the delayed beam at the entrance into the microscope. Inside the microscope, the light is reflected by a dichroic mirror that matches the excitation/emission wavelengths (FF545/650-Di01, Semrock, Rochester, NY, USA) and coupled into a 100 x, NA1.4 objective (Nikon, France). Emitted photons are then collected by the objective and focused on a pinhole of 150 μm. The emission photon stream is collimated and divided using a polarizing beamsplitter. In each created polarization channel, the photons are spectrally separated using dichroic mirrors (BS 649, Semrock, Rochester, NY, USA) and filtered using high quality emission filters (parallel: ET BP 585/65, Chroma, Bellows Falls, VT, USA and FF01-698/70-25, Semrock, Rochester, NY, USA, perpendicular: HQ 590/75 M, Chroma, Bellows Falls, VT, USA and FF01-698/70-25, Semrock, Rochester, NY, USA). Single photons are detected using Single Photon Avalanche Diodes. We use two MPD-1CTC (MPD, Bolzano, Italy) for the donor wavelength channels and two SPCM AQR-14 (Perkin Elmer, Fremont, CA, USA) for the acceptor wavelength channels. The output of the detectors is coupled into a TCSPC counting board (SPC-150, Becker&Hickl, Berlin, Germany), through a HRT41 router (B&H), using appropriate pulse inverters and attenuators. The sync signal that triggers the TCSPC board is provided by picking a small fraction of the light from the prompt path (reflected by a coverslip), and focusing it on an avalanche diode (APM-400, B&H).

### smFRET measurements

Measurements were performed at concentrations of 30-100 pM on SensoPlate 384 well plates (non-treated, Greiner Bio-One, France) passivated with 10mg/ml bovine serum albumin (BSA) in acquisition buffer with detergent for at least 1h prior to sample application. Samples were measured in acquisition buffer with detergent and ligand concentrations as indicated in the text. Measurements at saturating ligand concentration were carried out at 10 mM Glu, 100 μM LY37, 100 μM LY34, 1 mM LCCG-I, 1 mM DCG-IV and 1 mM LY35. Allosteric modulators BINA and Ro64 were supplemented at a final concentration of 10 μM. The effect of BINA at 500 nM was reversed by the addition of 10μM ro64. To study the effect of heterotrimeric human G_αi1_G_β1γ2_ on ECD reorientation 1 μM of the heterotrimer in the absence or presence of ligand was incubated with receptor (at approximately 30-100 pM) for 30 min at room temperature in the presence of 1 μM TCEP, 100 nM GDP, followed by the addition of 0.05 U/μl of apyrase (Sigma-Aldrich, France) and incubation for another 30min before acquisition.

### smFRET data analysis

Data analysis was performed using the Software Package for Multiparameter Fluorescence Spectroscopy, Full Correlation and Multiparameter Fluorescence Imaging developed in C.A.M. Seidel’s lab (http://www.mpc.uni-duesseldorf.de/seidel/) as described previously(Olofsson et al. 2014). A single-molecule event was defined as a burst containing of at least 40 photons with a maximum allowed interphoton time of 0.3 ms and a Lee-filter of 20. Photobleaching events were identified base on |TGX-TRR|<1 ms as described (Kudryavtsev et al. 2012). For some measurements, a minority of contaminating molecules with a long donor lifetime (>3 ns) were removed, as well as those with a lifetime >4ns in the acceptor channel for measurements in IGEPAL.

τ_DA_ vs E analysis and time windows analysis were performed using the PAM software(Schrimpf et al. 2018). The static FRET line for the τ_DA_ vs E analysis was plotted taking into consideration the excited state lifetime of the donor, and a 6 Å dye linker length.

Apparent FRET efficiencies (EPR), FRET efficiencies (E) and Stoichiometry were calculated using the conventions and recommendations made in (N. K. Lee et al. 2005) and (Hellenkamp et al. 2018) Precision, i.e.

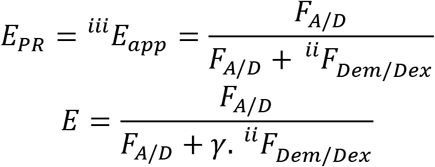

Where,

*^ii^F_xem/yex_* is the background corrected intensity in the X emission channel upon Y excitation. *F_A/D_* are the detected intensities in the acceptor emission channel upon donor excitation, corrected for background, donor leakage α (fraction of the donor emission into the acceptor detection channel) and direct excitation δ (fraction of the direct excitation of the acceptor by the donor-excitation laser) γ is the normalization factor that considers effective fluorescence quantum yields and detection efficiencies of the acceptor and donor.

### Additional Software

LRET data was analyzed using MARS (BMG Labtech) and displayed in GraphPad PRISM 7.05. FRET histograms were fitted and displayed using Origin 6 (Microcal Software, Inc.).

