## Supplementary Figures for "Allosteric modulators enhance agonist efficacy by increasing the residence time of a GPCR in the active state"

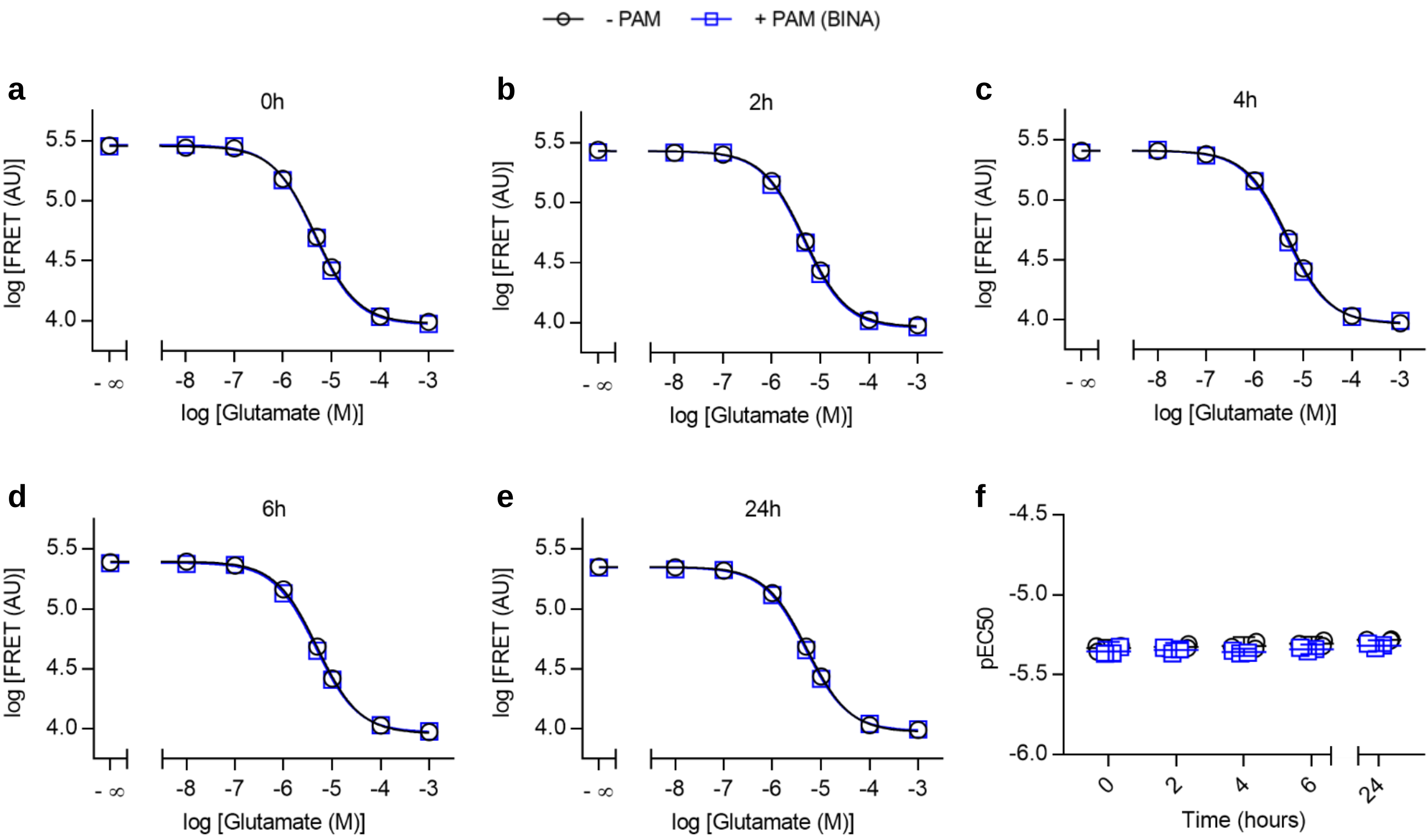


**Supplementary Figure 1: Functional assessment of mGluR2 in 0.05% IGEPAL by LRET.** a-e) Dose-response curves of Glutamate in the absence (Alone, black circles) and presence of 10µM BINA (+ BINA, blue squares) recorded after different time intervals of sample storage at room temperature. f) Change of pEC50 values over time, obtained from dose-response curves in a-e. Data are given as the mean of three biological replicates each measured in triplicate with errors given as standard deviation (a-e) or 95% confidence interval (f).

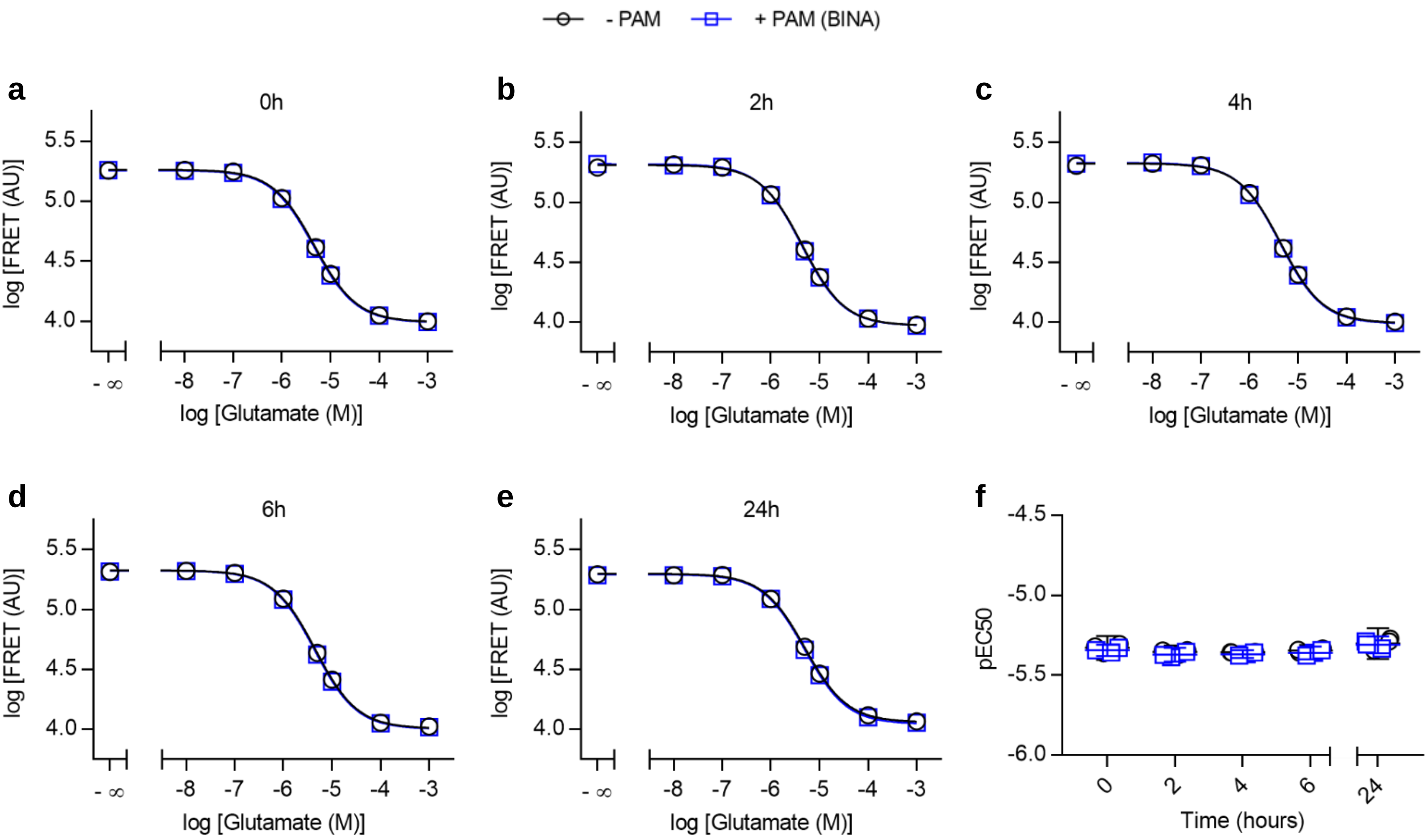


**Supplementary Figure 2: Functional assessment of mGluR2 in 0.05% DDM by LRET.** a-e) Dose-response curves of Glutamate in the absence (Alone, black circles) and presence of 10µM BINA (+ BINA, blue squares) recorded after different time intervals of sample storage at room temperature. f) Change of pEC50 values over time, obtained from dose-response curves in a-e. Data are given as the mean of three biological replicates each measured in triplicate with errors given as standard deviation (a-e) or 95% confidence interval (f).

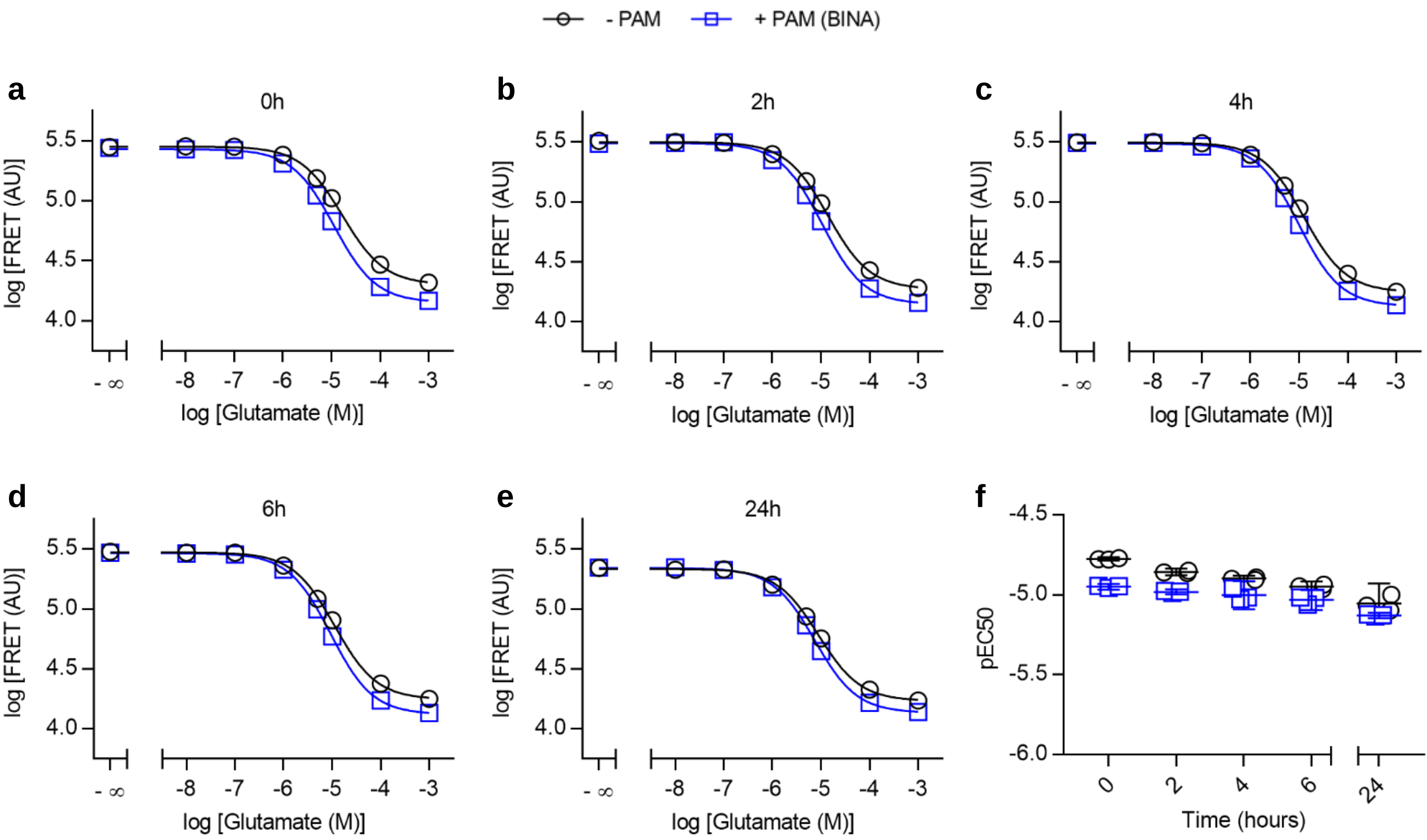


**Supplementary Figure 3: Functional assessment of mGluR2 in 0.05% DDM + 0.008% CHS by LRET.** a-e) Dose-response curves of Glutamate in the absence (Alone, black circles) and presence of 10µM BINA (+ BINA, blue squares) recorded after different time intervals of sample storage at room temperature. f) Change of pEC50 values over time, obtained from dose-response curves in a-e. Data are given as the mean of three biological replicates each measured in triplicate with errors given as standard deviation (a-e) or 95% confidence interval (f).

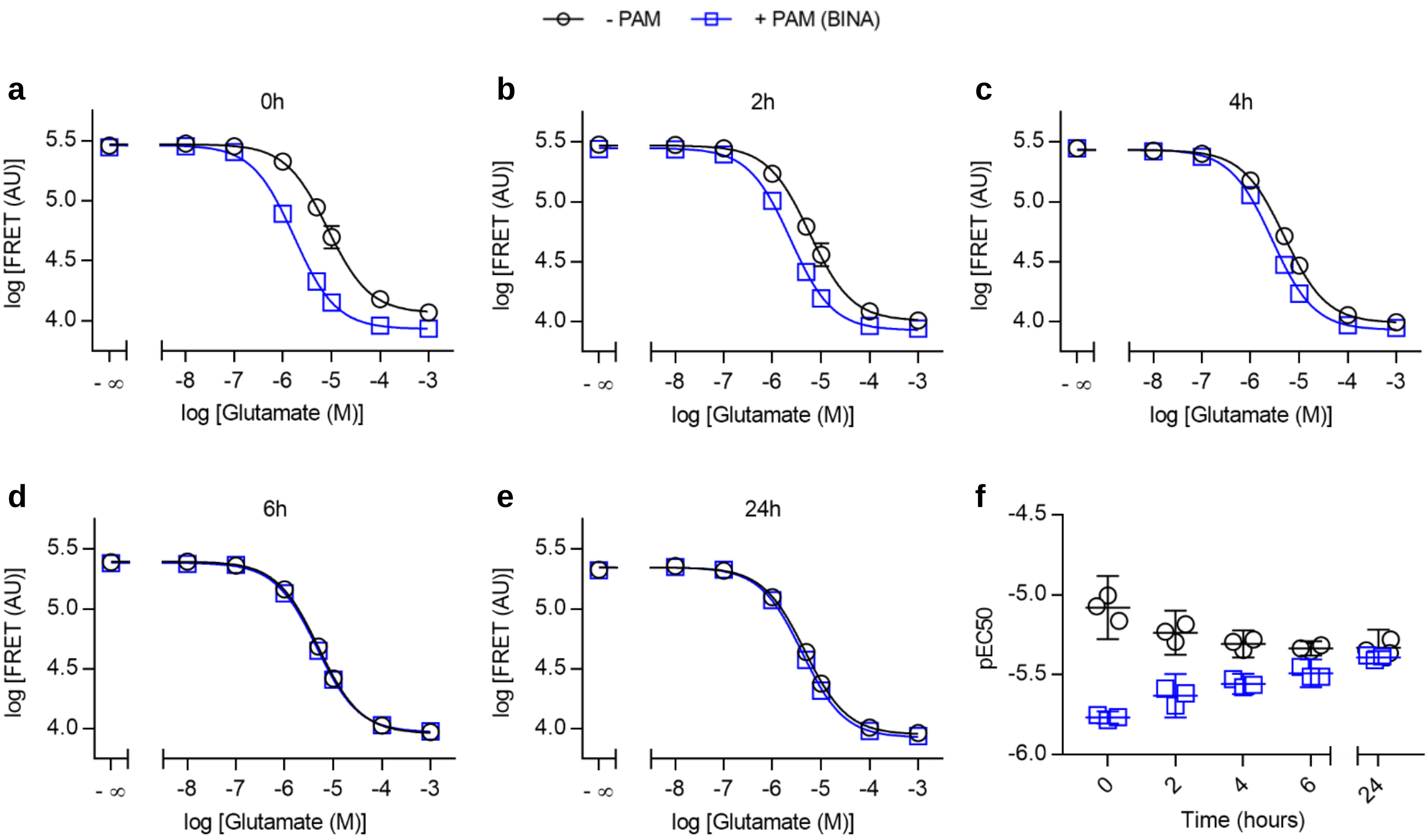


**Supplementary Figure 4: Functional assessment of mGluR2 in 0.005% LMNG by LRET.** a-e) Dose-response curves of Glutamate in the absence (Alone, black circles) and presence of 10µM BINA (+ BINA, blue squares) recorded after different time intervals of sample storage at room temperature. f) Change of pEC50 values over time, obtained from dose-response curves in a-e. Data are given as the mean of three biological replicates each measured in triplicate with errors given as standard deviation (a-e) or 95% confidence interval (f).

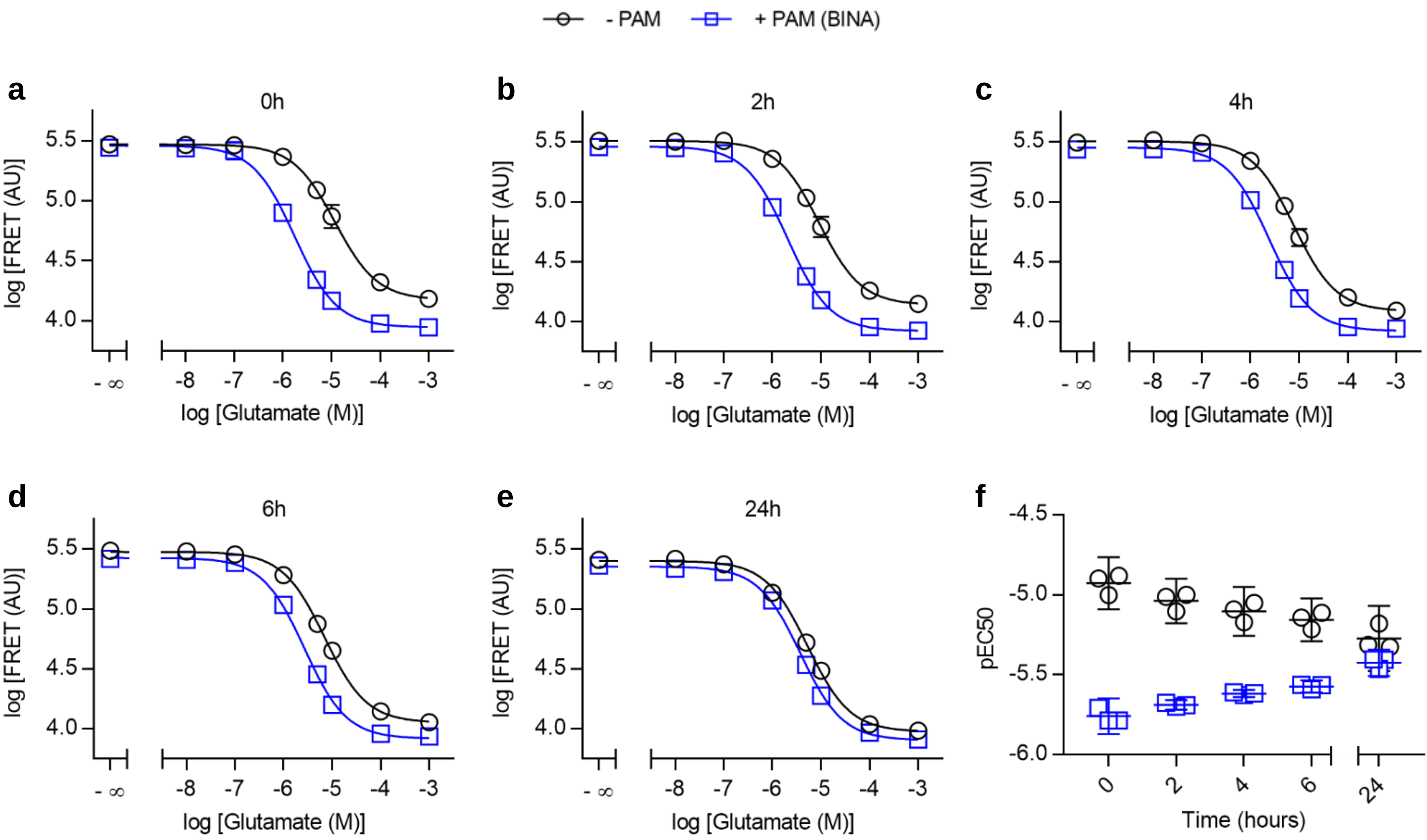


**Supplementary Figure 5: Functional assessment of mGluR2 in 0.005% LMNG + 0.0002% CHS by LRET.** a-e) Dose-response curves of Glutamate in the absence (Alone, black circles) and presence of 10µM BINA (+ BINA, blue squares) recorded after different time intervals of sample storage at room temperature. f) Change of pEC50 values over time, obtained from dose-response curves in a-e. Data are given as the mean of three biological replicates each measured in triplicate with errors given as standard deviation (a-e) or 95% confidence interval (f).

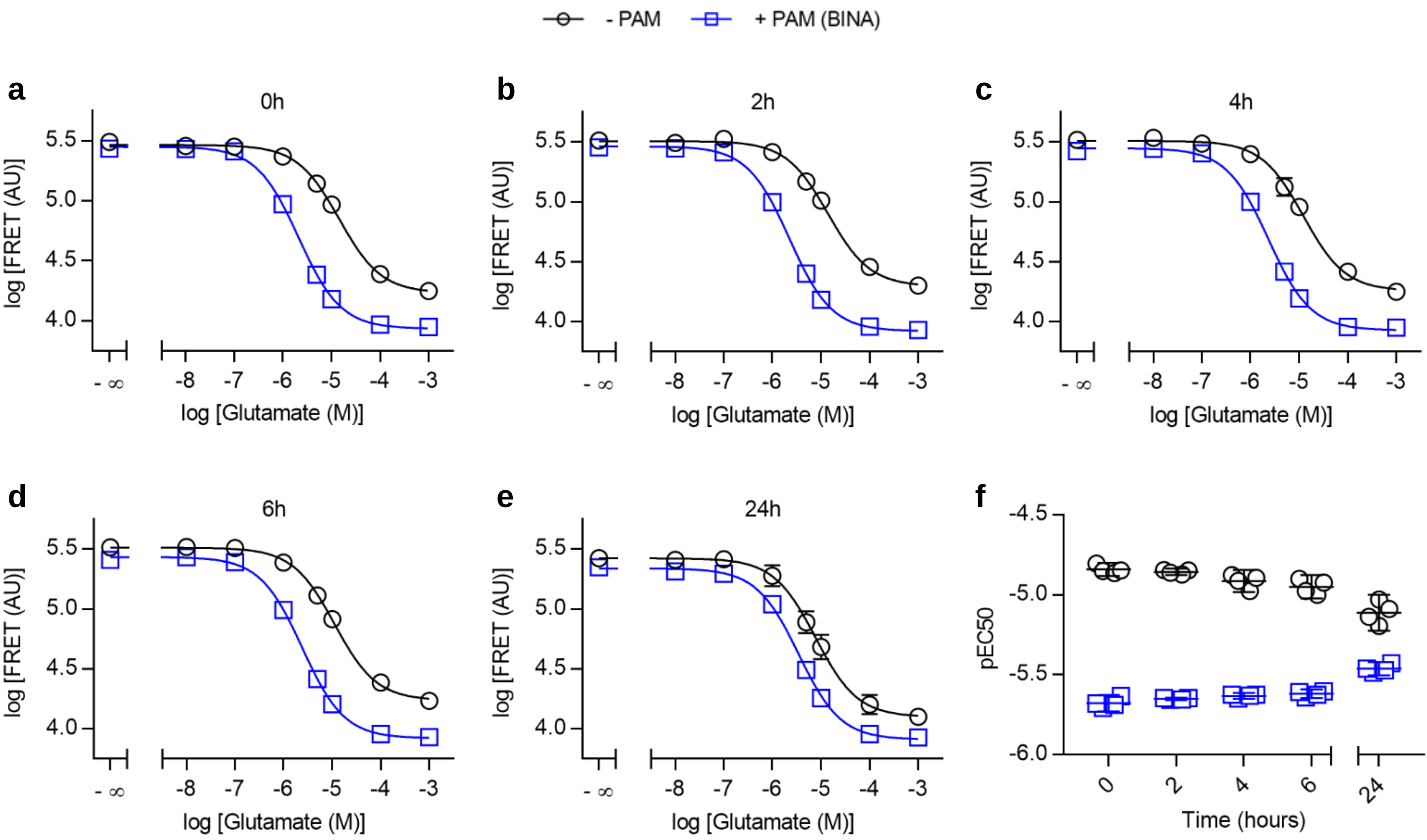


**Supplementary Figure 6: Functional assessment of mGluR2 in 0.005% LMNG + 0.0004% CHS by LRET.** a-e) Dose-response curves of Glutamate in the absence (Alone, black circles) and presence of 10µM BINA (+ BINA, blue squares) recorded after different time intervals of sample storage at room temperature. f) Change of pEC50 values over time, obtained from dose-response curves in a-e. Data are given as the mean of three biological replicates each measured in triplicate with errors given as standard deviation (a-e) or 95% confidence interval (f).

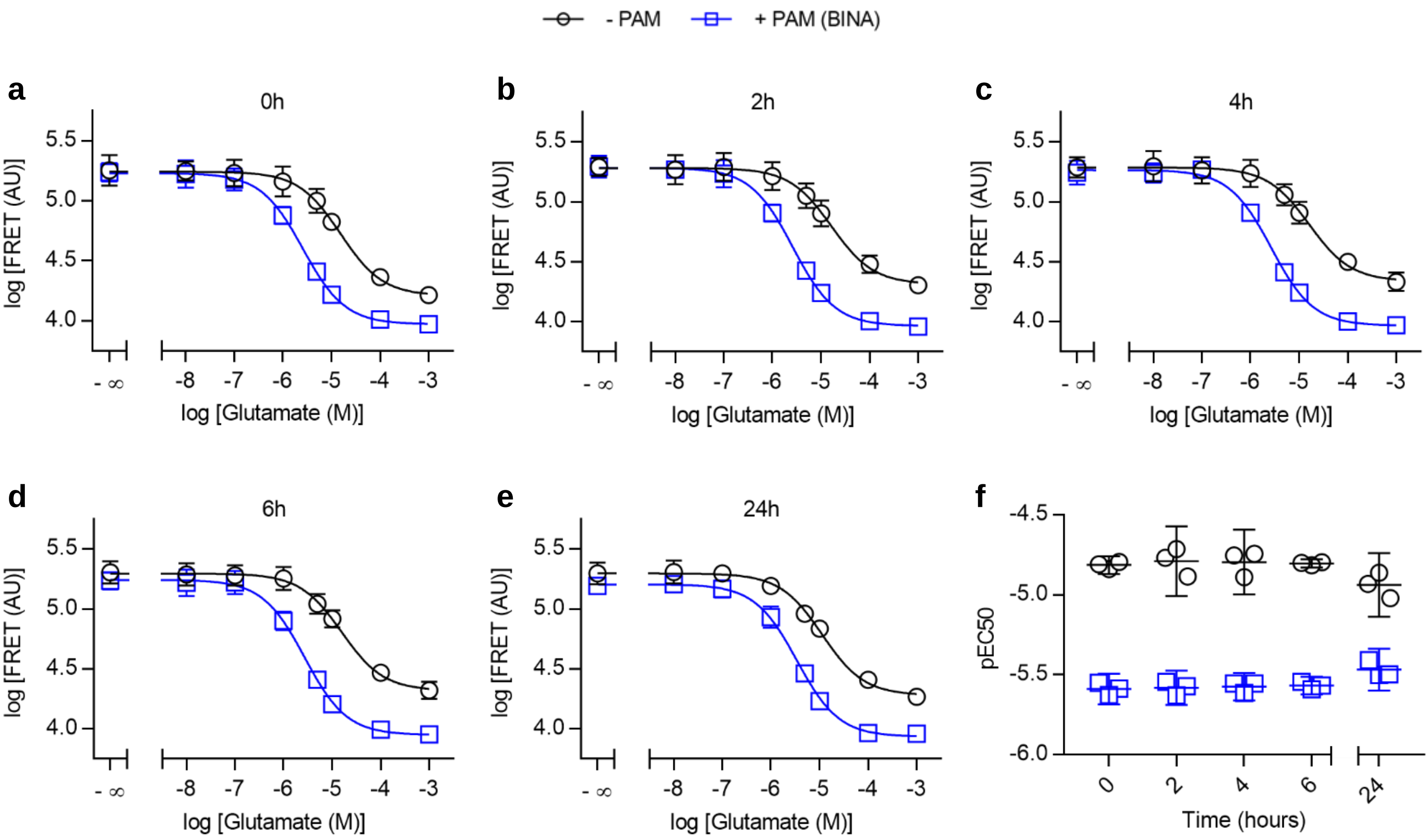


**Supplementary Figure 7: Functional assessment of mGluR2 in 0.005% LMNG + 0.0008% CHS by LRET.** a-e) Dose-response curves of Glutamate in the absence (Alone, black circles) and presence of 10µM BINA (+ BINA, blue squares) recorded after different time intervals of sample storage at room temperature. f) Change of pEC50 values over time, obtained from dose-response curves in a-e. Data are given as the mean of three biological replicates each measured in triplicate with errors given as standard deviation (a-e) or 95% confidence interval (f).

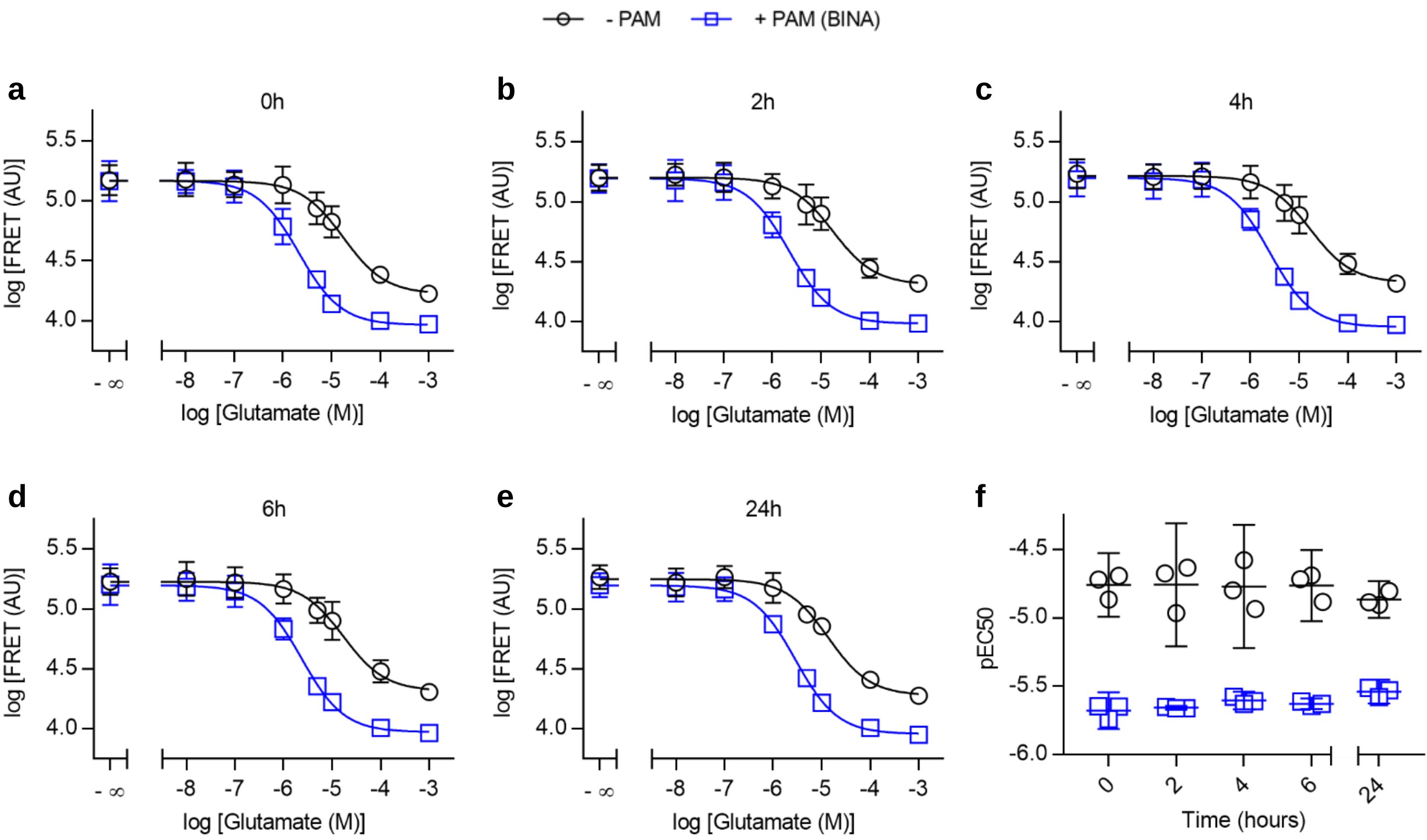


**Supplementary Figure 8: Functional assessment of mGluR2 in 0.005% LMNG + 0.0004% CHS + 0.0025% GDN by LRET.** a-e) Dose-response curves of Glutamate in the absence (Alone, black circles) and presence of 10µM BINA (+ BINA, blue squares) recorded after different time intervals of sample storage at room temperature. f) Change of pEC50 values over time, obtained from dose-response curves in a-e. Data are given as the mean of three biological replicates each measured in triplicate with errors given as standard deviation (a-e) or 95% confidence interval (f).

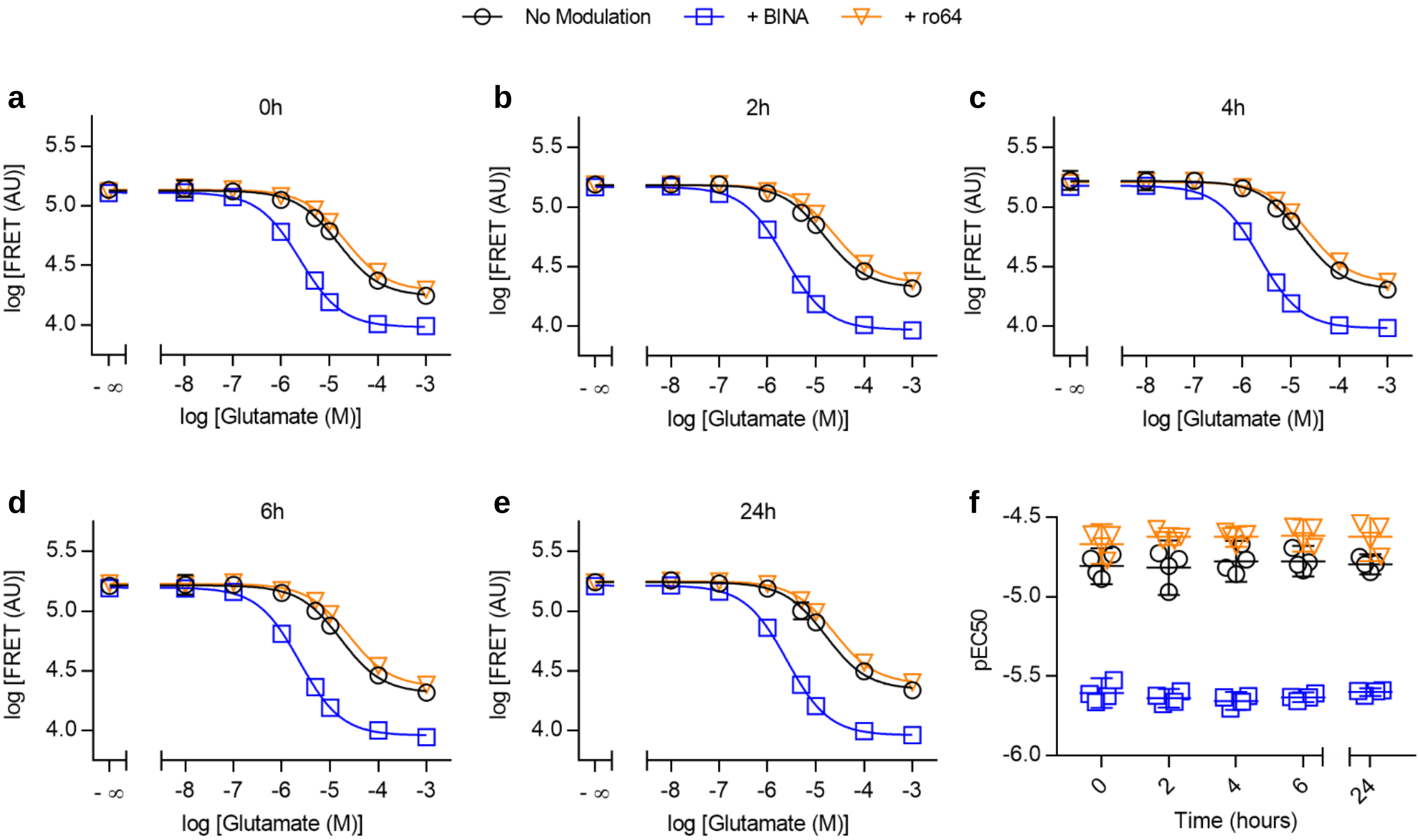


**Supplementary Figure 9: Functional assessment of mGluR2 in 0.005% LMNG + 0.0004% CHS + 0.005% GDN by LRET.** a-e) Dose-response curves of Glutamate in the absence (Alone, black circles) and presence of 10µM BINA (+ BINA, blue squares) and 10µM ro64 (+ ro64, orange triangles) recorded after different time intervals of sample storage at room temperature. f) Change of pEC50 values over time, obtained from dose-response curves in a-e. Data are given as the mean of three biological replicates each measured in triplicate with errors given as standard deviation (a-e) or 95% confidence interval (f).

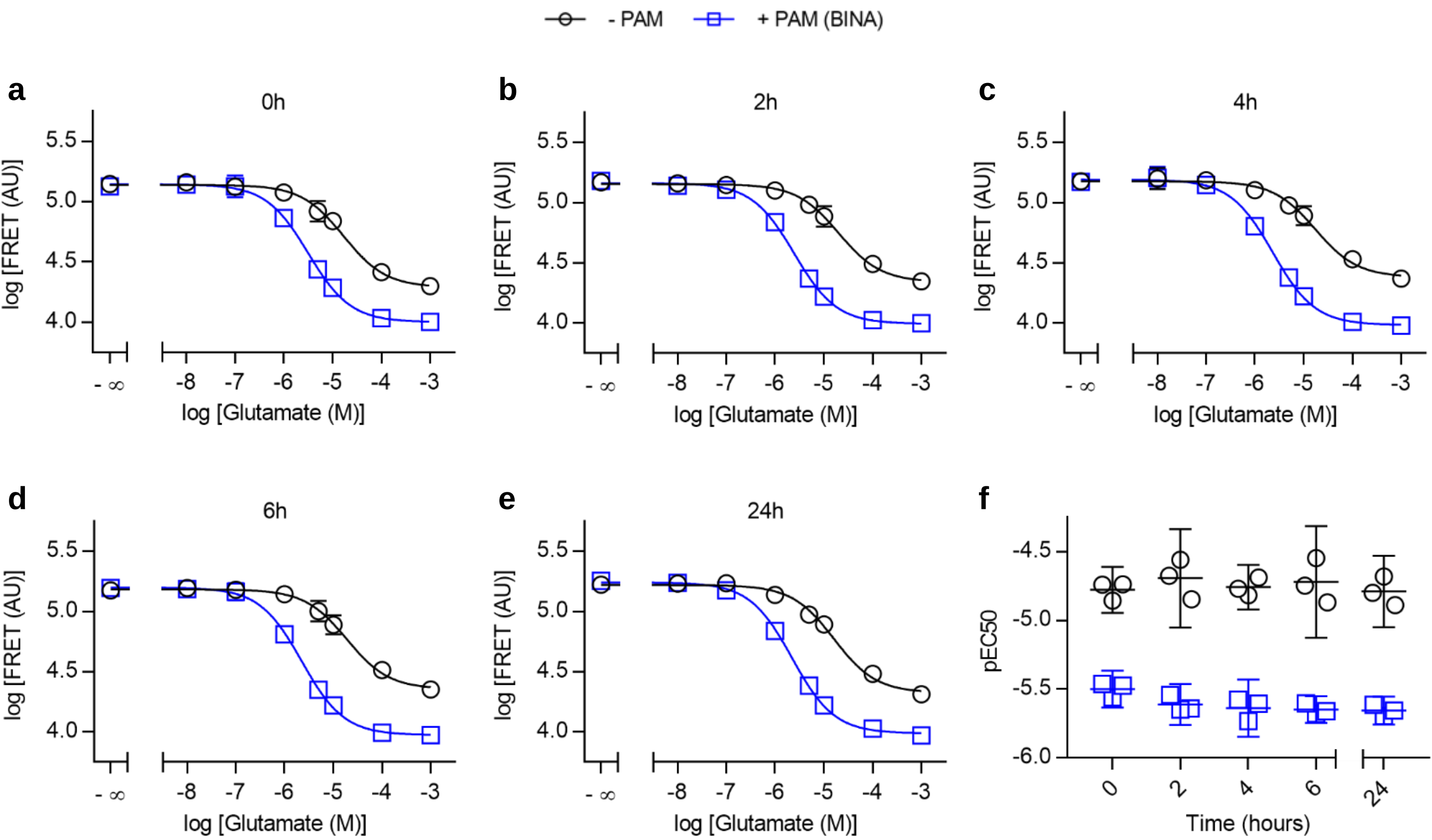


**Supplementary Figure 10: Functional assessment of mGluR2 in 0.005% LMNG + 0.0004% CHS + 0.01% GDN by LRET.** a-e) Dose-response curves of Glutamate in the absence (Alone, black circles) and presence of 10µM BINA (+ BINA, blue squares) recorded after different time intervals of sample storage at room temperature. f) Change of pEC50 values over time, obtained from dose-response curves in a-e. Data are given as the mean of three biological replicates each measured in triplicate with errors given as standard deviation (a-e) or 95% confidence interval (f).

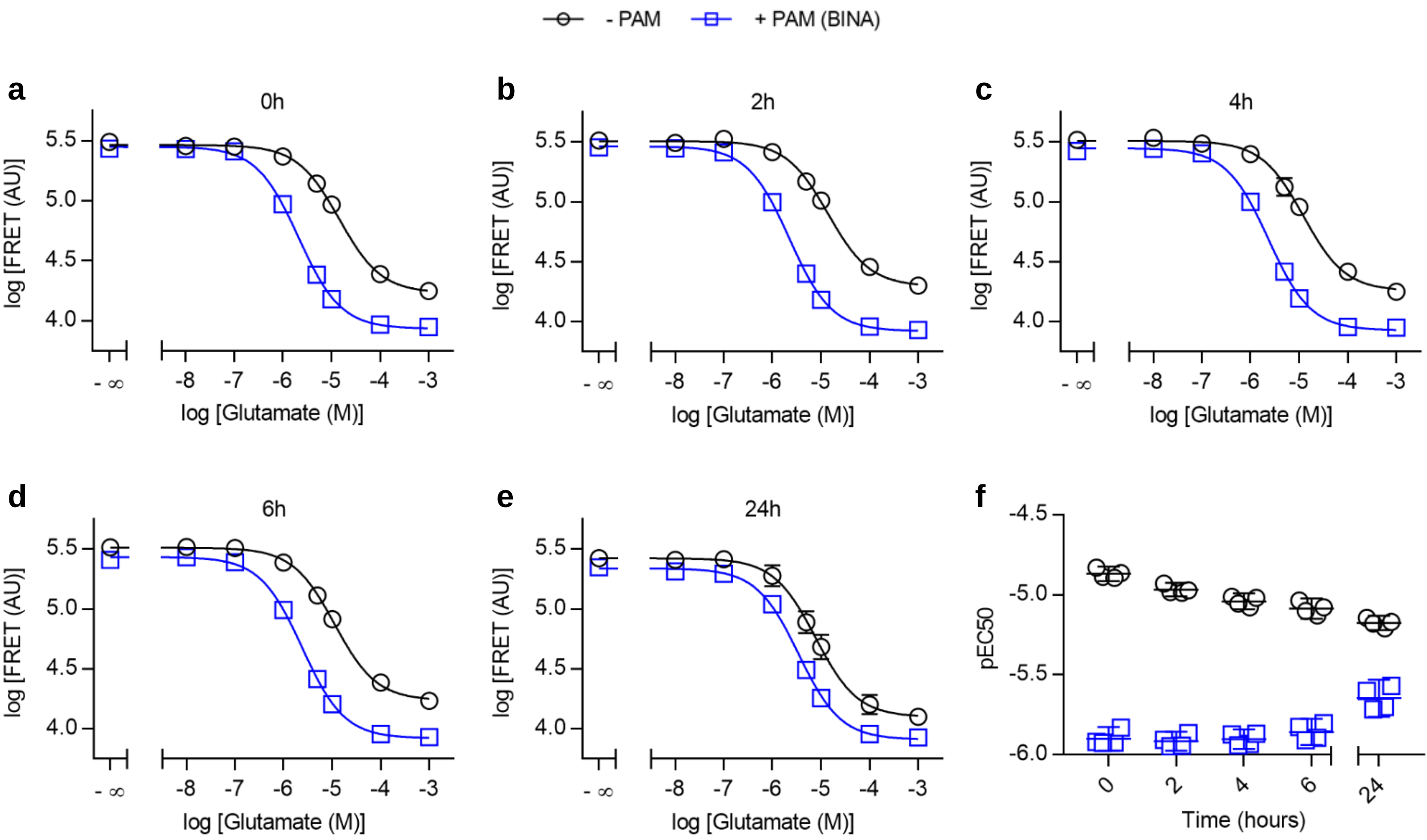


**Supplementary Figure 11: Functional assessment of mGluR2 in 0.005% LMNG + 0.005% GDN by LRET.** a-e) Dose-response curves of Glutamate in the absence (Alone, black circles) and presence of 10µM BINA (+ BINA, blue squares) recorded after different time intervals of sample storage at room temperature. f) Change of pEC50 values over time, obtained from dose-response curves in a-e. Data are given as the mean of three biological replicates each measured in triplicate with errors given as standard deviation (a-e) or 95% confidence interval (f).

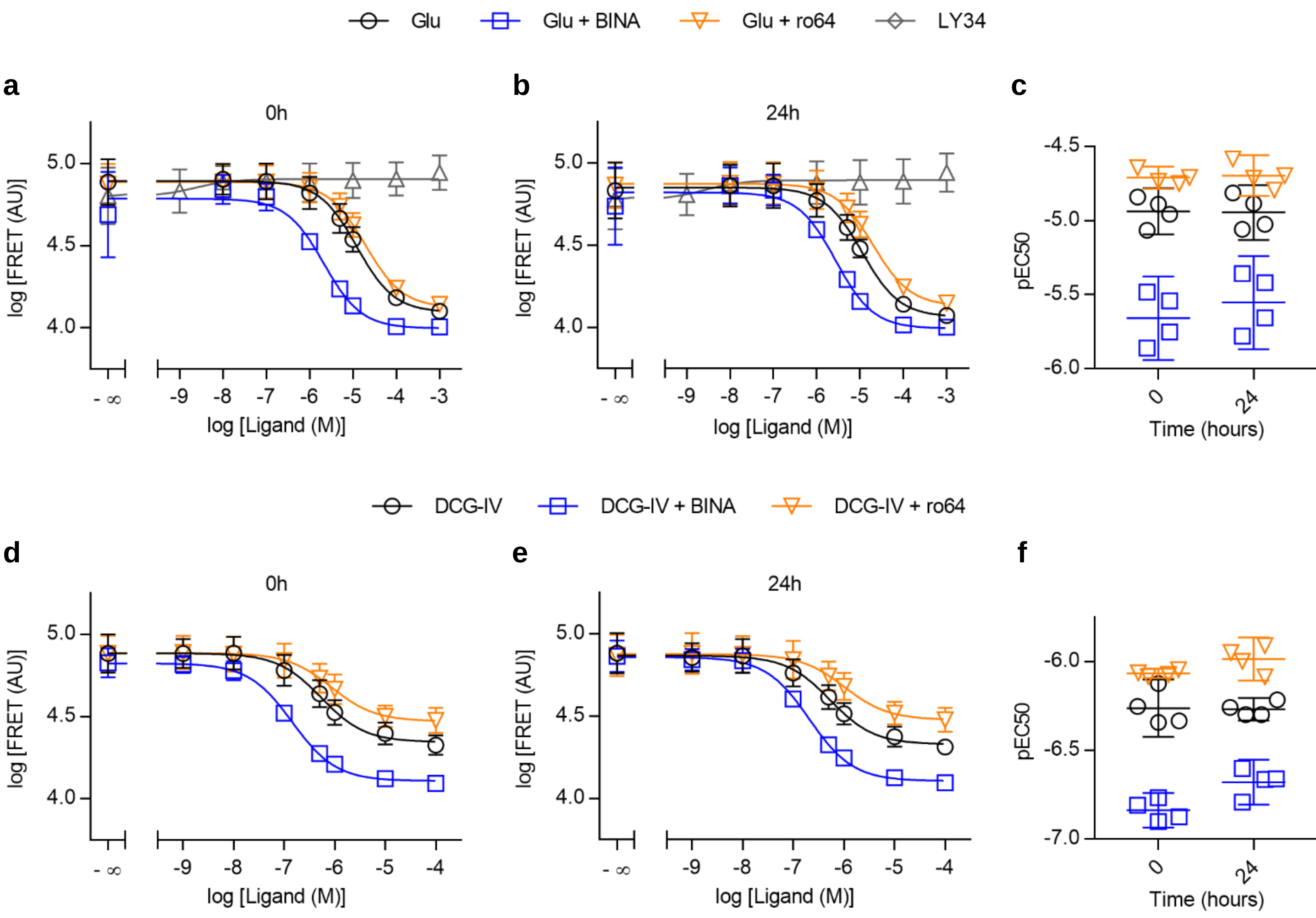


**Supplementary Figure 12: Orthosteric ligand response of mGluR2 in membranes by LRET.** a-b) Dose-response curves of LY34 alone (grey triangles) and Glu in the absence (black circles) and presence of 10µM BINA (blue squares) and 10µM ro64 (orange triangles) recorded after different time intervals of sample storage at room temperature. c) Change of pEC50 values over time, obtained from dose-response curves in a-b. d-e) Dose-response curves of DCG-IV in the absence (black circles) and presence of 10µM BINA (blue squares) and 10µM ro64 (orange triangles) recorded after different time intervals of sample storage at room temperature. f) Change of pEC50 values over time, obtained from dose-response curves in d-e. Data are given as the mean of three biological replicates each measured in triplicate with errors given as standard deviation (a-e) or 95% confidence interval (f).

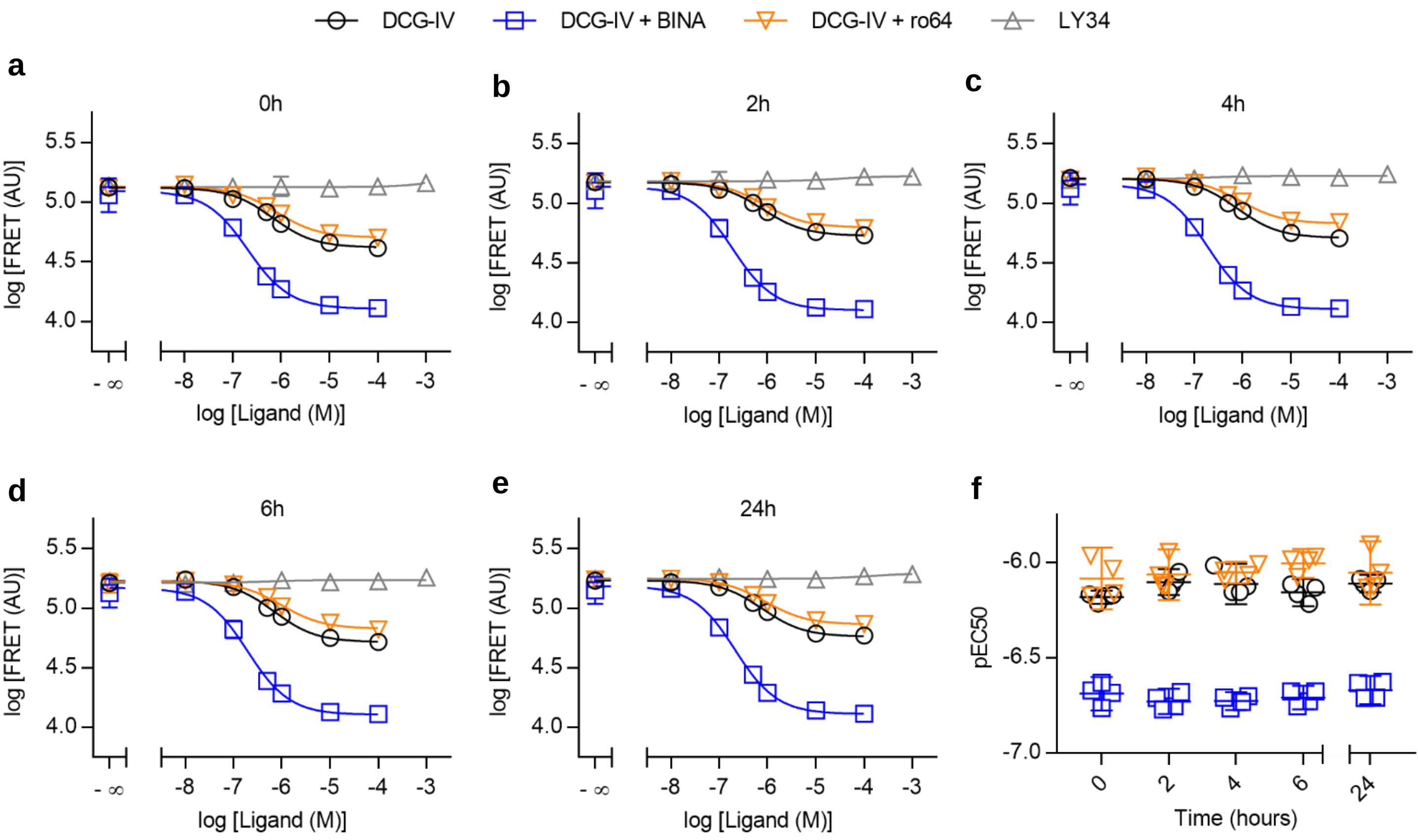


**Supplementary Figure 13: Orthosteric ligand response of mGluR2 in 0.005% LMNG + 0.0004% CHS + 0.005% GDN by LRET.** a-e) Dose-response curves of LY34 alone (grey triangles) and DCG-IV in the absence (black circles) and presence of 10µM BINA (blue squares) and 10µM ro64 (orange triangles) recorded after different time intervals of sample storage at room temperature. f) Change of pEC50 values over time, obtained from dose-response curves in a-e. Data are given as the mean of three biological replicates each measured in triplicate with errors given as standard deviation (a-e) or 95% confidence interval (f).

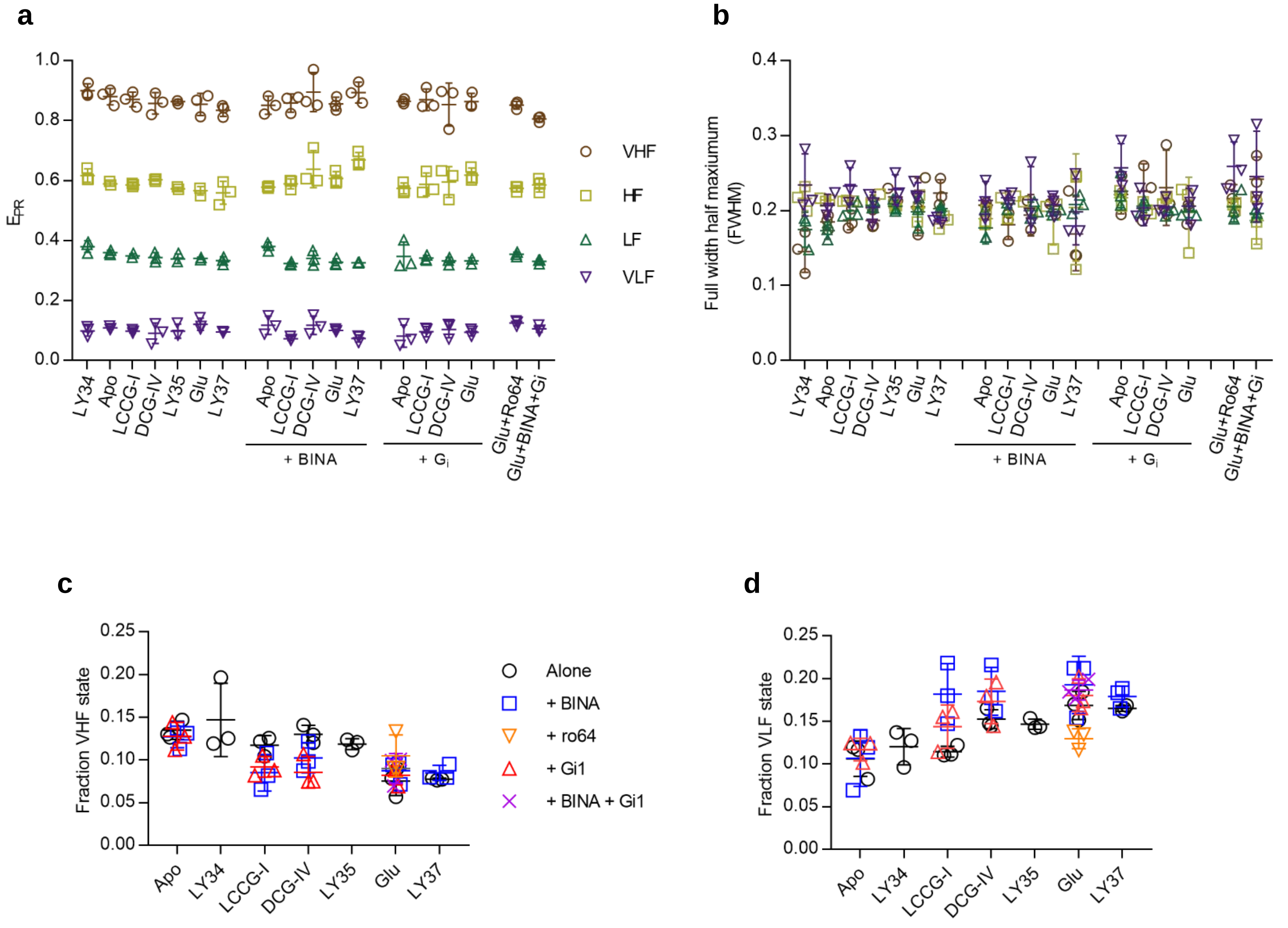


**Supplementary Figure 14 : Determination of average $E_{PR}$ and FWHM and fraction of VHF and VLF.** a-b) FRET distributions were fitted with four gaussians alternatingly keeping either $E_{PR}$ or FWHM variable until no further improvement was achieved. c-d) Fraction of VHF and VLF at fixed mean $E_{PR}$ and FWHM at various ligand conditions. Data are given as the mean of three biological replicates with errors given as standard deviation.

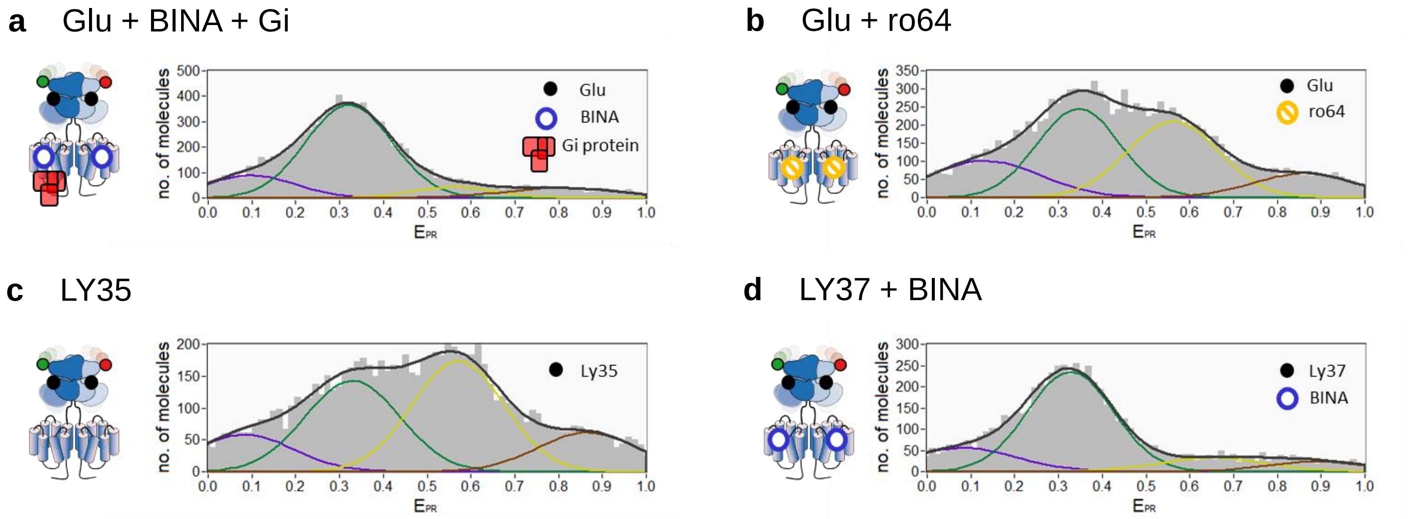


**Supplementary Figure 15: FRET histograms for additional ligands.** Representative $E_{PR}$ histograms with Glu+BINA+Gi (a), Glu+ro64 (b), LY35 (c) and LY37+BINA (d). FRET distributions were fitted with four gaussians (black curves) alternatingly keeping either $E_{PR}$ or FWHM variable until no further improvement was achieved. The sum of the four gaussians is displayed in red.

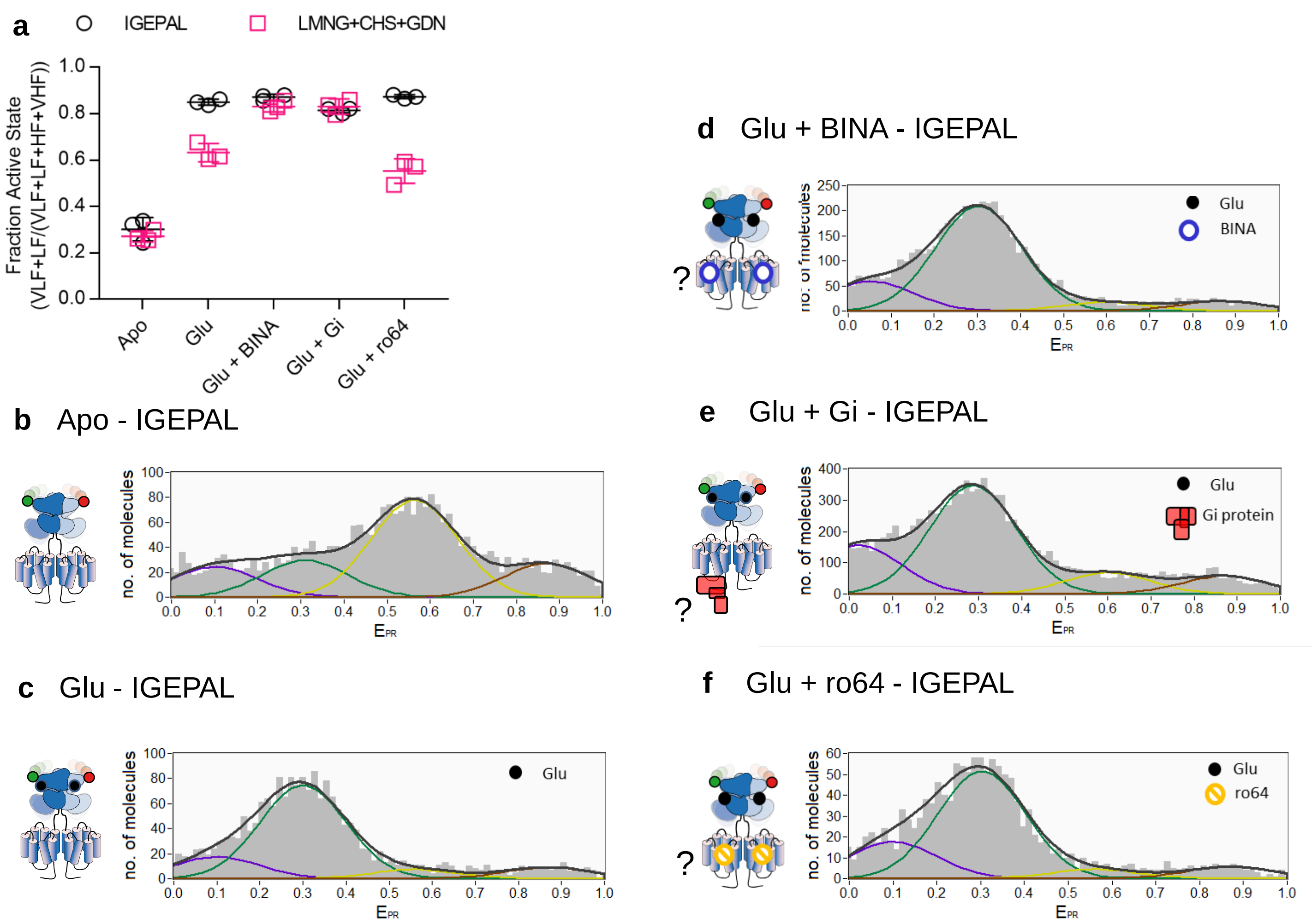


**Supplementary Figure 16:** Comparison of mGluR2 activation in IGEPAL and LMNG-CHS-GDN micelles. a) Comparison of the degree of activation in response to different ligands shows that in IGEPAL micelles Glu already leads to a maximal VFT reorientation, while BINA, $G_i$ and ro64 exhibit no effect. b-f) Corresponding histograms of DA species, where contaminant species at long lifetimes are removed, fitted with fixed FWHM (0.2) and VHF (0.87) but variable VLF, LF and HF. As no effect of the histogram are observed, the binding of BINA, Gi and ro64 to mGluR2 in IGEPAL cannot be confirmed.

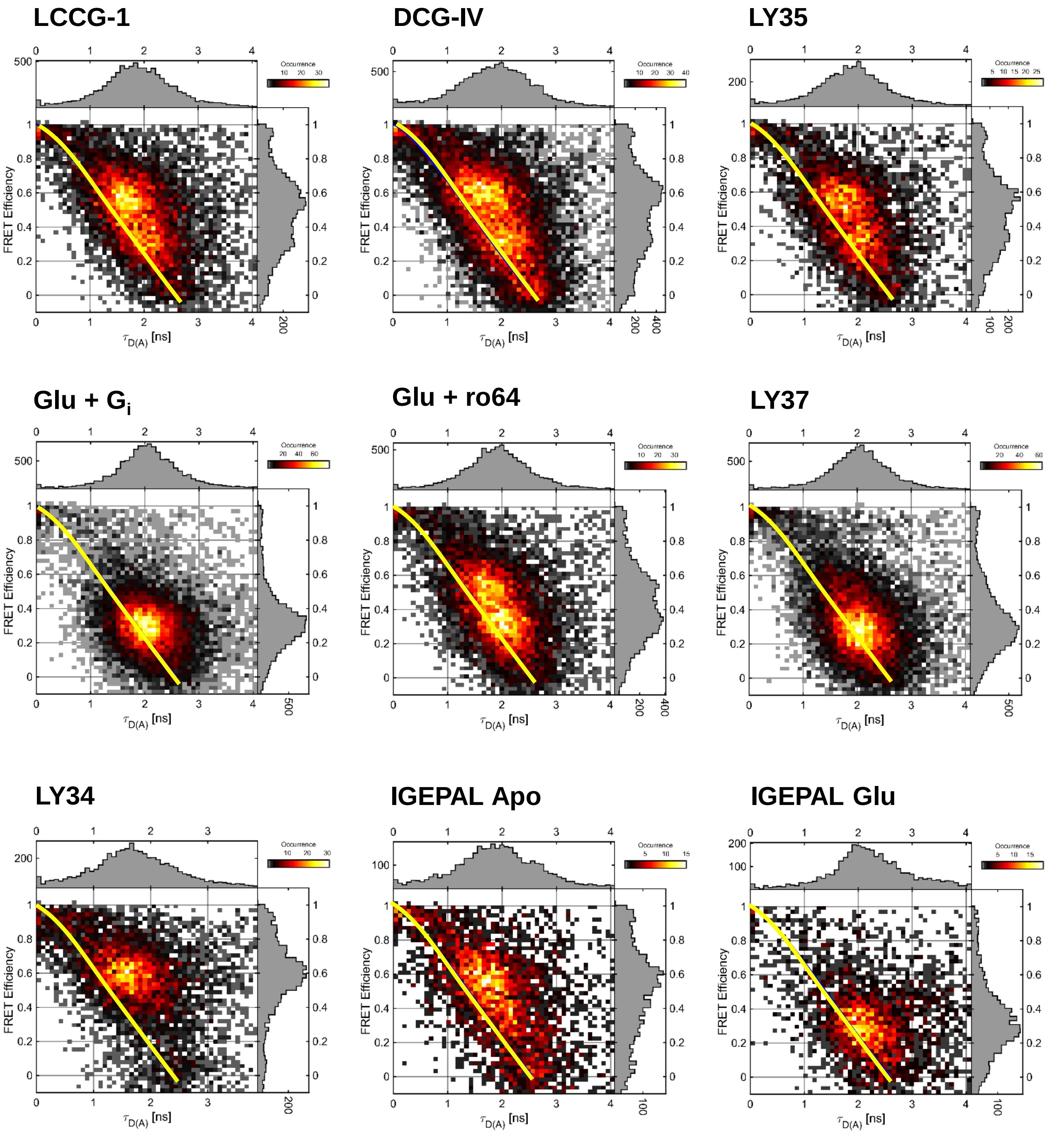


**Supplementary Figure 17: $\tau_{DA}$ vs *E* histogram under additional conditions.** Representative $\tau_{DA}$ vs E histograms for mGluR2 dimers solubilized with LMNG-CHS-GDN. Top row) in the presence of partial agonists (LCCG-1, DCGIV, LY35, respectively), center row) Glu+Gi, Glu+ro64 and superagonist LY37, bottom row, left) antagonist LY34. Bottom row center and right) Dimers in IGEPAL in the absence (Apo) and presence of Glu. The yellow line represents the “static FRET” line.

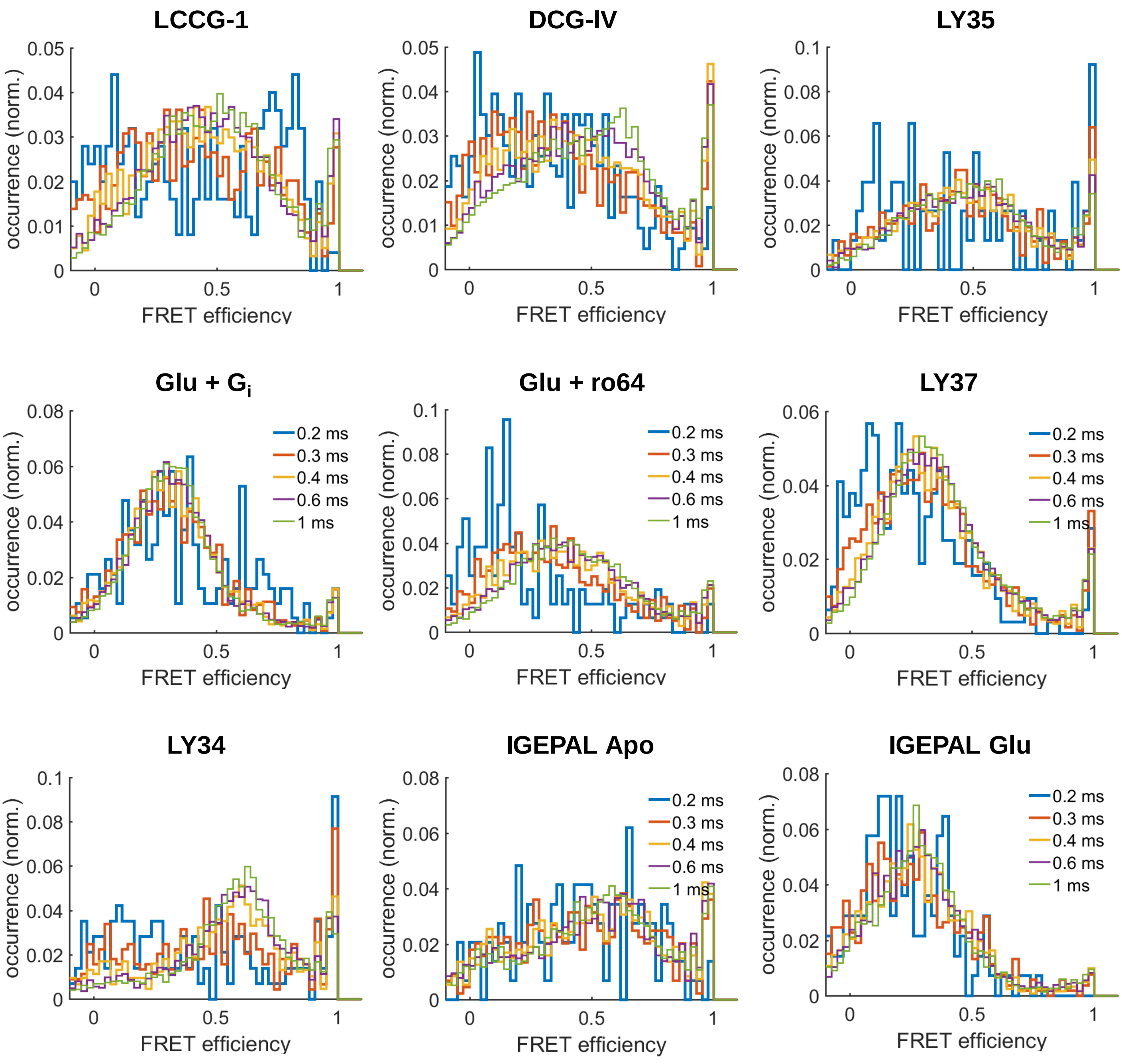


**Supplementary Figure 18: Time window analysis under additional conditions.** Representative TWA histograms at 0.2ms to 1ms integration times for mGluR2 dimers. Top row) in the presence of partial agonists (LCCG-1, DCGIV, Ly35). Center row) in the presence of Glu+Gi, Glu+ro64 and LY37. Bottom row) In the presence of antagonist LY34, in IGEPAL micelles in the absence (Apo) and presence of Glu.
